# Allosteric Gating Mechanism Regulates Odorant Selectivity and Antagonism in Odorant Receptors

**DOI:** 10.64898/2026.07.06.736896

**Authors:** Ning Ma, Mona Marie, Dan Takase, Clàudia Llinàs del Torrent Masachs, Jeremy Aguilar, Aashish Manglik, Hiroaki Matsunami, Nagarajan Vaidehi

## Abstract

Odorant receptors (ORs) belong to class A G protein coupled receptors that detect diverse small molecules, yet the steps that link odorant association to receptor mediated selectivity remains incompletely defined. Here we combined 1.26 milliseconds of all-atom odorant association Molecular Dynamics simulations with Markov state modeling and cell-based cAMP measurements to examine two human ORs receptors that recognize chemically distinct odorants. In the class I receptor OR51E2, propionate associates via two extracellular pathways gated for selectivity by residues in the extracellular loop 2 and 3. The longer alkyl chain heptanoate occupies this gate and reduces propionate association and signaling which is consistent with the observed antagonist behavior of heptanoate. Pocket-expanding mutations at F155 and L158 allosterically regulate ECL2-ECL3 gate by permitting longer-chain fatty acids to adopt fully inserted poses that support gate closure, while also attenuating propionate responses. In the class II receptor OR1A1, hydrophobic odorants partition into the membrane and reach the orthosteric site mostly through multiple transmembrane paths. A mutational scan of gating residues and analysis of intermediate-state occupancies suggest that an orthosteric substitution at G108 can allosterically bias odorant association path choice. Together, these results support a model in which odorant association paths and gate residence, together with allosteric coupling between the orthosteric site and entry gates, contribute to odorant specificity and antagonism in odorant receptors.

## Introduction

Odorant receptors (ORs) belong to class A G protein coupled Receptor (GPCR) family of transmembrane (TM) proteins. Each OR is tuned to recognize a specific set of odorants, and collectively the OR repertoire is capable of binding a vast array of diverse odorants.^1,2^ Although ORs are structurally homologous to class A GPCRs, they exhibit differences in their structural and functional characteristics.^3–5^ For example, since ORs respond to a broad array of odorants they show sequence and structural diversity in the residues lining their binding pockets.^6^ Rather than relying on highly specific, tightly constrained binding interactions, ORs may employ more flexible recognition mechanisms based on broader structural complementarity.^7–9^ These observations raise the possibility that ORs may utilize distinct odorant recognition mechanisms compared to other class A GPCRs.

Structural characterization of ORs remains challenging due to their low expression levels, inherent instability when purified, and conformational flexibility, resulting in limited high-resolution structural data compared to other GPCR subfamilies.^10,11^ Recent emergence of three-dimensional structures of both class I ORs that get activated by hydrophilic odorants and class II ORs that get activated by hydrophobic odorants have provided the first look at the odorant binding sites in ORs.^4,5,7,12–14^ The three-dimensional structure of human OR51E2 in complex with propionate (abbreviated as C3 henceforth) has shown that the receptor’s selectivity to short chain fatty acids come from the small size of the binding pocket . We have shown that mutation of residues L158^4.57^ and F155^4.60^ located in the bottom of the propionate binding site to alanine expands the binding pocket leading to activation of the mutant OR51E2 by longer chain fatty acid like heptanoate (abbreviated as C7 henceforth).^4^ Using molecular dynamics (MD) simulations we showed that the closure of extracellular loop2 (ECL2) with ECL3 is an important feature of OR51E2 activation. Breaking of R262^6.59^-Q181^ECL2^ interaction, a critical interaction responsible for ECL2-ECL3 closure, will eliminate OR51E2 activity. However, although the binding pocket is enlarged in F155A and L158A, the shorter-chain propionate activates OR51E2 less strongly than in wild type (WT). This shows that other structural factors besides the sufficient size of the odorant binding site could be responsible for odorant recognition by OR51E2. Three dimensional structures of engineered class II ORs such as consOR1, consOR2 and consOR4 showed that odorant binding site alone is not sufficient to explain odorant selectivity.^5^

Previous extensive MD simulations of odorant association process in class A GPCRs have shown the presence of extracellular residues that confer odorant selectivity to receptors with identical odorant binding sites.^15–19^ Our previous work mapped the association paths of the small odorant N,N-dimethylcyclohexylamine into the olfactory receptor mTAAR7f which belongs to non-OR GPCRs but responds to odorous amines, and identified an entry gate formed by ECL2 and ECL3 that permits ligand access to the binding site.^20^ Despite these advances, fundamental questions remain regarding the precise molecular mechanisms that govern odorant entry pathways and the coordinated roles of multiple extracellular loops in determining odorant selectivity by ORs. The dynamic involvement of EC loops and/or involvement of lipid bilayer in odorant association, the determinants of odorant binding, and the mechanistic basis for how orthosteric mutations allosterically modulate gate dynamics remain poorly understood. Additionally, there is very limited information on the odorant association mechanisms of hydrophobic odorants in class II ORs. Traditional experimental approaches, while valuable for characterizing functional responses, lack the temporal and spatial resolution necessary to capture the transient intermediate states and pathway-dependent effects that likely underlie olfactory receptor discrimination. These mechanistic gaps represent a significant barrier to understanding how ORs achieve their remarkable ability to distinguish between structurally similar molecules while maintaining the promiscuity necessary for broad chemical detection.

Here we advance understanding of mechanisms of odorant selectivity by both OR51E2, (class I OR) and OR1A1 (class II OR), using 1.26 milliseconds of odorant association MD simulations combined with Markov State Modeling (MSM) to identify recurrent association states and generate a mechanistic model that regulate odorant selectivity. Our unbiased odorant association MD simulations sampled associated poses consistent with odorant-bound structures. Our results show that the hydrophilic short-chain fatty acid enter the OR51E2 binding site through the EC gate formed by ECL2 and ECL3, similar to mTAAR7f. On the other hand, hydrophobic aromatic odorants dissolve into the lipid bilayer and enter the OR1A1 binding site through the TM helices via multiple association pathways. We predicted through these simulations that heptanoate (C7) acts as an antagonist in the presence of propionate (C3) in the wild type OR51E2 that was tested in multi-odorants mixture experiments. Such unprecedented insights advance our understanding of odorant selectivity by ORs.

## Results

### Two entry paths and one ECL gate mediate propionate (C3) association in OR51E2 (a Class I OR)

We performed brute force all-atom C3 association MD simulations of the OR51E2 (Apo form by removing odorant from PDB ID: 8F76)^4^ embedded within a POPC lipid bilayer with cholesterol, with near Emax concentrations of C3 in solution (See Supplementary Table 1 and Extended Methods for justifications and details). We conducted five independent MD simulations each 15 μs long adding up to an aggregate of 75 μs (Fig. 1A, left panel). Multiple complete association events of C3 were observed (Fig. 1A, right panel, Supplementary Table 2), enabling identification of recurrent association paths and ligand-position intermediates. Given the finite-sampling uncertainty observed in MSM validation (Supplementary Data and Extended Method), we interpret the resulting path assignments qualitatively rather than as converged statistical estimates of association-path probabilities. We used the MSM analysis with distances between odorants and residues in the odorant-binding site (OBS) as reaction coordinates. Through this approach, we aimed to identify key binding events and estimate the occupancies of critical meta stable states, shedding light on the molecular basis of OR odorant selectivity.

**Figure 1.**
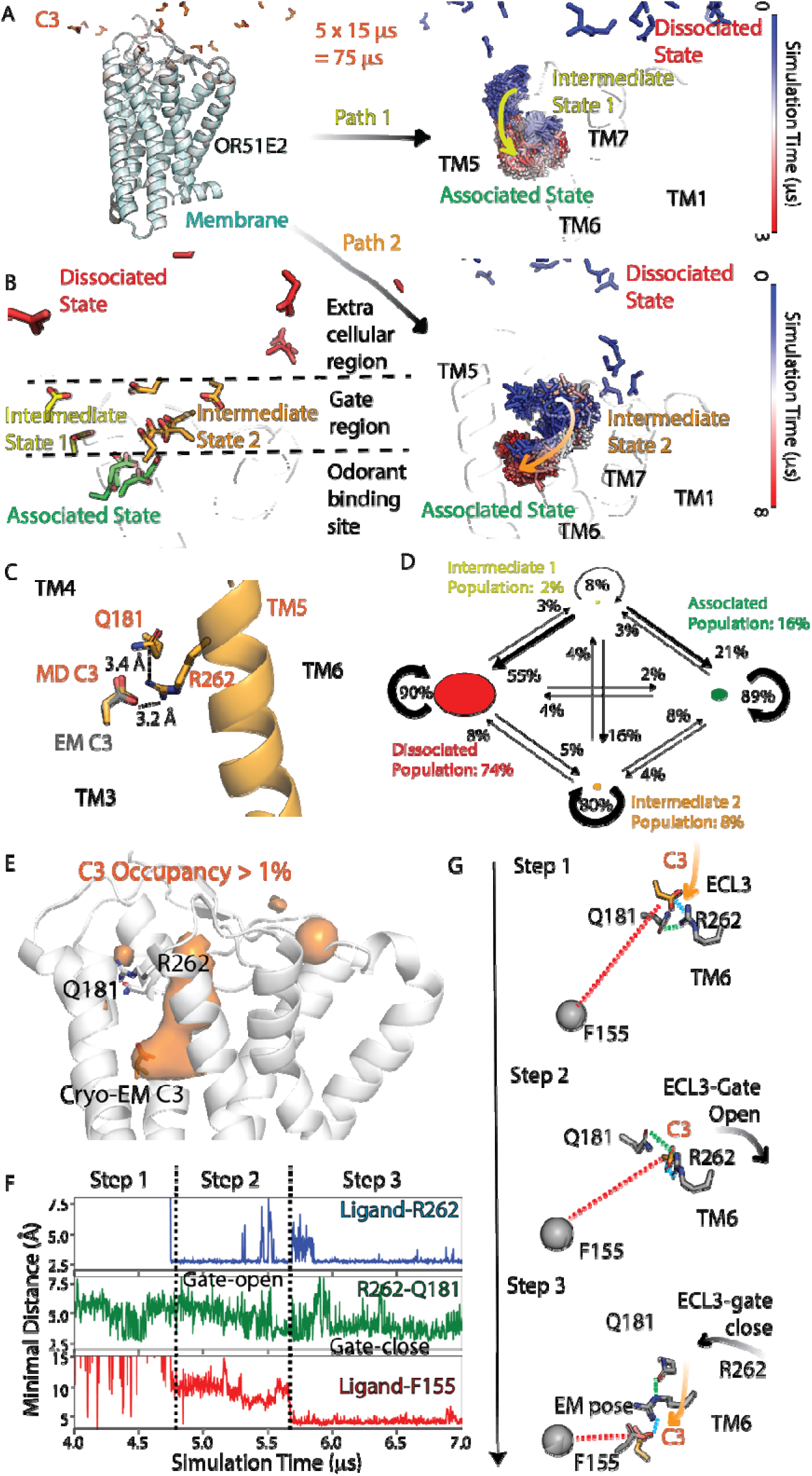
C3 association with the orthosteric odorant-binding site of OR51E2. (A) System setup for odorant association MD simulations. OR51E2 shown as cartoon without C3 bound was embedded in POPC/Cholesterol bilayer, with C3 molecules randomly distributed in the solution. Path 1 (top right panel) for C3 association begins with C3 entering the receptor through the ECL2-ECL3-TM5 gate. The odorant conformations, sampled every 20 ns, is depicted as sticks colored by simulation time. Path 2 (bottom right panel) involves C3 entering through the ECL2-ECL3-TM7 gate. (B) Representative odorant conformations from the 14 micro states derived from MSM overlaid on OR51E2 structure, and odorants are colored according to their associated (green)/ intermediate (light green and brown)/ dissociated states (red) (The three odorant states are categorized by the distance between odorant and odorant binding site, which is separated by black dash lines into three regions: extra cellular regions, gate regions, and odorant binding site. See Supplementary Methods for details for definition of these states). Multiple odorants from dissociated states are shown, only one odorant is shown from each of the other states. (C) Representative C3 conformation from the associated state overlaid with cryo-EM odorant binding pose (pdb ID: 8f76). The distance between R262-C3, and R262-Q181 is labeled by black dash lines. (D) A MSM state transition network showing the transition probabilities (on the arrows) between four states. The population of each state is indicated using corresponding color-coded numbers. The number within the curved arrows shows the probability of transitions within that macrostate (E) C3 occupancy from OR51E2-C3 simulations, the orange cloud shows the structural regions where C3 occupancy is greater than 1%. (F) Time series plots of the minimal distance between C3-R262, R262-Q181 and C3-F155. (G) Representative structures showing the process of C3 association in three steps. In step1 of the C3 association pathway, C3 interacts with R262 while R262 still interacts with Q181. Step 2, R262-Q181 gate opens, R262 facilitates the transfer of C3 into the binding site. Step 3, C3 is transferred into binding site, R262-Q181 closed afterwards.

The MSM analysis of OR51E2-C3 simulations revealed 14 distinct microstates (See Extended Methods for details). We overlaid the representative odorant conformations from these 14 microstates on the OR51E2 structure and defined the associated state (odorants in OBS), dissociated state (odorants in water), and intermediate states (odorants positioned on the paths between water and OBS), based on the location of C3 in OR51E2 (Fig. 1B). A representative of the odorant associated state of C3 overlays with the cryo-EM structure of C3 bound to OR51E2 with 0.5 Å root mean square deviation (RMSD) in the odorant atom coordinates, supporting the relevance of our C3 association MD simulations (Fig. 1C).

Two distinct, non-overlapping odorant association paths were identified by comparing the intermediate state conformations and the original odorant association trajectories, (Fig. 1A right panels). The first path involves odorant entry from the extracellular solution phase through an opening at the ECL2–ECL3–TM5 interface (path 1), any odorant conformations located along this path is considered as intermediate state 1. While the second pathway originates similarly in the extracellular solution but proceeds through an opening at the ECL2–ECL3–TM7 interface (path 2), with odorant conformations along this path considered as intermediate state 2. Notably, path 2 resembles the odorant association path previously identified in mTAAR7f.^20^

MSM results also reveal that the two paths differ in the probability of odorant entry (Fig. 1D). In our MSM analysis, we use the distances between C3 and residues in its binding site as reaction coordinates (see Methods). We use the population of a given metastable state (intermediate state in this case, see method for detail) to estimate the fraction of time the system is expected to occupy that state under equilibrium conditions. The two C3 association paths differed in their apparent residence behavior near the extracellular gate. Direct trajectory inspection suggested that path 1 more directly connected the extracellular region to the orthosteric pocket, whereas path 2 involved broader sampling of intermediate gate-associated positions (Fig. 1D). Consistent with this view, odorant occupancy maps, which report the spatial density of odorant atoms around the receptor, showed a more localized C3 occupancy along path 1 and a wider occupancy cloud along path 2 (Fig. 1E; Supplementary Table 2). These results support a qualitative model in which path 1 represents a more direct association path and path 2 represents a longer-lived intermediate sampling path. Because MSM validation indicated finite-sampling uncertainty, we do not interpret the MSM-derived population or transition-probability differences as quantitative kinetic estimates. Our results so far indicate that odorant association proceeds with distinct dynamics along these two pathways.

### A Three-Step Mechanism of Odorant Association in OR51E2

We analyzed the different steps in the association pathways of C3. In Step 1, the initial event of odorant association in the EC loop region involves the formation of an ionic interaction between the carboxylic acid group of C3 and the side chain of R262 (Fig. 1F blue curve, and 1G step1 panel). This interaction is maintained across intermediate and associated states. Experimentally, mutation of R262 to alanine abolishes C3-induced activation of OR51E2,^4^ highlighting its critical role in odorant binding .

In Step 2, the hydrogen bond between R262 (in ECL3) and Q181 (in ECL2) is broken to allow C3 to transfer into the orthosteric pocket (Fig. 1F green curve, and 1G step 2 panel). The R262-Q181 separation coincided with formation of the C3-R262 interaction. The movement of ECL3 away from ECL2 creates a transient opening that permits odorant entry, effectively functioning as a dynamic “gate”. When ECL2 and ECL3 remain in close contact, the gate remains closed and prevents odorant access to the binding site. After C3 enters the binding site, R262 acts as a gatekeeper by re-forming the hydrogen bond with Q181 to stabilize the gate-closed conformation which we speculate leads to receptor activation. Interestingly, in the dissociated state, the R262–Q181 interaction fluctuates between gate-open and gate-closed conformations, potentially facilitating initial odorant recognition by R262. Disruption of this interaction marks, or is associated with, a transition to the intermediate state for odorant association, in which the odorant is located between solution and OBS (Fig. 1B).

In Step 3, full odorant association is marked by a decrease in the distance between C3 and F155, from ∼10 Å to ∼5 Å, reflecting the odorant entry into the orthosteric binding site (Fig. 1F red curve, and 1G step 3 panel). Residue F155, located at the base of the orthosteric site within the TM domain, forms direct contact with C3 in the cryo-EM structure and serves as a structural reference point for monitoring odorant association. Once the odorant is fully associated, the ECL2–ECL3 gate closes, as evidenced by the re-formation of the R262–Q181 interaction (Fig. 1C, Fig. 1F, green curve in step 3).

Taken together, our results highlight the importance of the ECL2–ECL3 gate and identify R262 as a key determinant of odorant recognition that helps capture and retain the odorant from solution.

### C7 antagonizes OR51E2 by occupying the ECL2-ECL3 gate and stabilizing an inactive gate-open conformation

Our previous work showed that C7 does not activate OR51E2 WT (Supplementary Fig. 1A).^4^ The above proposed three-step association led to a testable hypothesis that odorants that complete steps 1 and 2 but fail at step 3 which might achieve partial entry but not proper closure of the ECL2-ECL3 gate, and might act as an antagonist by occupying the gate region.

This hypothesis can be tested by simulating the association of C7 by itself as well as in mixture with C3. Cryo-EM structures of OR51E2 revealed a compact odorant-binding site (volume ∼31 Å³) precisely shaped to accommodate small size odorant. Computational docking of C7 into Cryo-EM structure suggested severe steric clashes with OBS residues, predicting C7 would be unable to bind. F155A and L158A mutations, which remove these bulky side chains and expand the binding pocket, enable C7 to robustly activate the receptor (Supplementary Figs. 1B,1C). Surprisingly, these same mutations simultaneously diminish C3 activity despite the larger odorant binding site volume. This observation raises two critical questions: If C7 cannot fit in the wild-type pocket, does it simply fail to bind, or does it interact with the receptor in a non-productive manner? And what is the mechanistic basis for this size-dependent selectivity switch? To answer these questions, we performed five MD runs (each run 15 μs long) of Emax concentration of C7 association to WT OR51E2 and the two F155A, L158A OR51E2 mutants (See Supplementary Table 1 and Extended Methods for justifications and details), then performed similar MSM analysis as previously.

Similar to the association of C3, C7 also associates with OR51E2 via two paths (Figs. 2A, 2B, 2C). Path 1 of C7 differs slightly from that of C3 in that it initially inserts its hydrophobic tail into the lipid bilayer and subsequently drifts into the odorant binding site through the ECL2–ECL3–TM5 opening (Fig. 2A, Supplementary Fig. 2A). In contrast, path 2 of C7 is similar to that observed for C3 (Fig. 2B). For C7 association in the F155A and L158A mutants, both path 1 and path 2 were observed (Supplementary Figs. 2B-2E). To cross compare the intermediates from these pathways, we pooled intermediates from all paths into a single metastable intermediate state and used its population as a single descriptor (Supplementary Fig. 7F).

**Figure 2.**
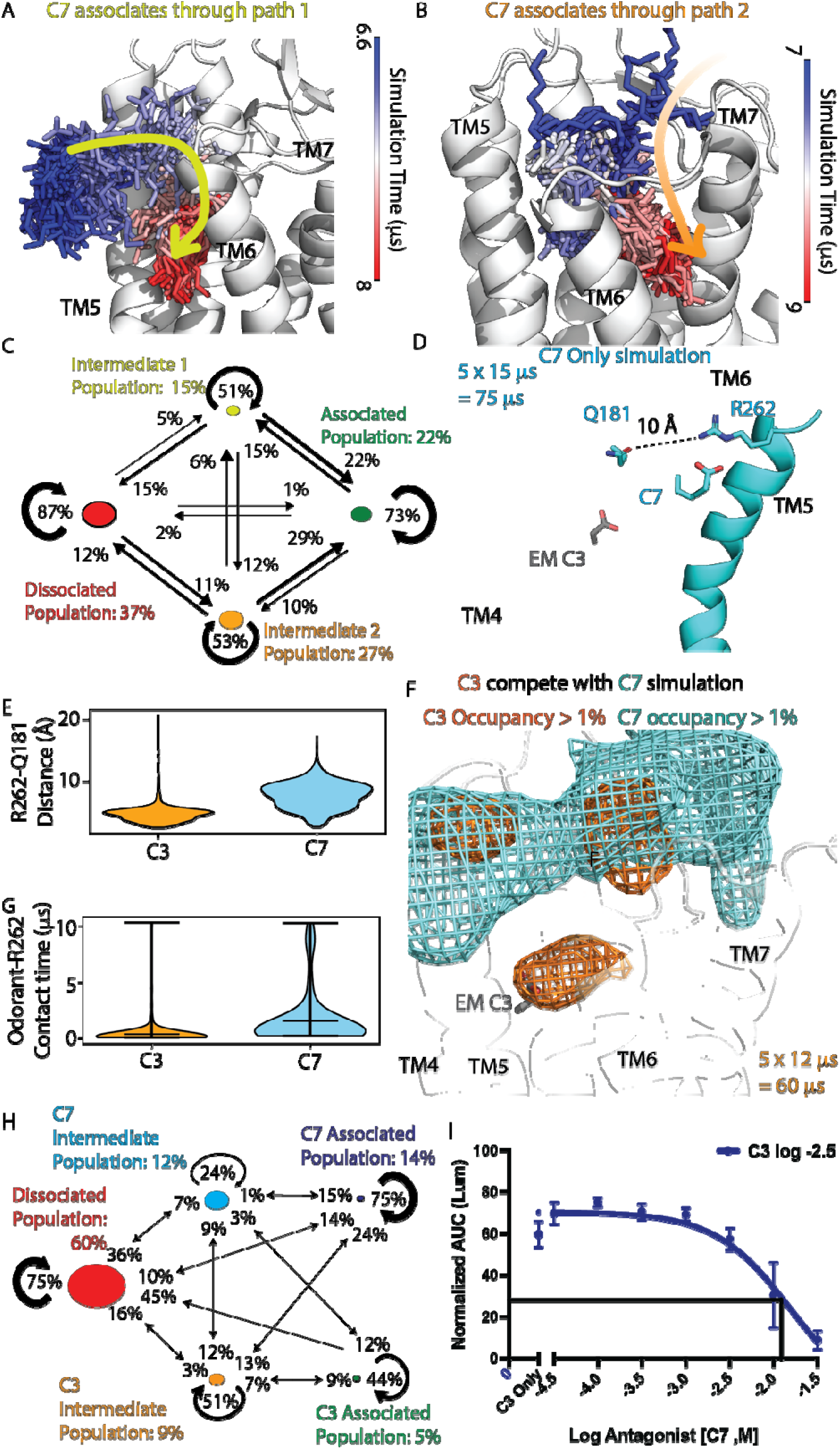
C7 is an antagonist of OR51E2. (A,B) Two paths for C7 association identified from MSM analysis of the MD simulation trajectories. (C) MSM state transition probability network illustrating the C7 association process. (D) Representative structure of C7 associated states, with binding pose of cryo-EM C3 shown. (E) Comparison of the Q181-R262 distance distributions in C3 and C7 association simulations. (F) Structural regions with greater than 1% occupancy density of C3 atoms (orange) and C7 atoms (cyan) from the C3-C7 mixture simulations. (G) Comparison of persistence of contact between odorant-R262 for C3 and C7 (percentage of MD snapshots showing the contact). (H) MSM state transition network illustrating the C3-C7 mixture simulation. (I) Experimental normalized AUC curve showing the effect of titrating C7 into C3-OR51E2 mixture.

A key difference between C3 and C7 association simulations lies in the final associated conformations of C7. The compact volume of the OBS in WT OR51E2 does not fully accommodate C7. As a result of C7 association, its hydrophobic tail inserts into the binding site while its head group remains solvent-exposed, forming a contact with R262 and maintaining the ECL2–ECL3 gate in an open conformation (Fig. 2D). Proper R262-Q181 interaction is impeded in such conformation, suggesting a possible non-productive association state (Fig. 2E).

In contrast, the expanded volume of the OBS introduced by F155A or L158A mutations enables full accommodation of C7, with both its tail and head group inserted into the binding pocket (Supplementary Figs. 2F, 2G). This accommodation allows the proper ECL2–ECL3 gate-closed conformation, representing the productive association state like that of C3 in WT OR51E2. We also simulated the association of C6 and C8 alkyl acids in the two OR51E2 mutants (10 runs each 5μs long). Similarly, odorant association simulations of C8 in F155A OR51E2 mutant and C6 in L158A OR51E2 mutant show complete odorant association and stable ECL2–ECL3 gate-closed conformation (Supplementary Figs. 2H, 2I, 13 and 10 association events were observed in C8-F155A and C6-L158A simulation, respectively, as shown in Supplementary Table 2).

Also, C7 exhibits a substantially higher intermediate population than C3 (Supplementary Fig. 7F). Because C7 and C3 access OR51E2 through the same paths, the greater population of C7 in intermediate states could hinder C3 from entering. Together with C7’s tendency to adopt non-productive conformations upon association, we hypothesis that C7 acts as an antagonist of C3 at WT OR51E2.

To examine a potential mechanism by which C7 antagonizes C3 responses, we performed simulations of WT OR51E2 with both C3 and C7 at near Emax concentration in same simulation runs (5 runs each run 12 μs long), allowing direct competition between the two odorants. Odorant occupancy density calculations show that C7 occupies the ECL2–ECL3 gate region more frequently than C3, as evidenced by a wider odorant occupancy distribution (Fig. 2F), higher C7-R262 contact residence time (Fig. 2G), and MSM-derived apparent occupancies showed a similar trend, with C7 modestly enriched in intermediate states relative to C3 (Fig. 2H). Notably, in the presence of C7, the MSM-derived apparent occupancy of C3-associated states was lower in the mixture simulations than in the C3-only simulations. (Fig. 2H, Supplementary Fig. 7F). Also, C3 showed reduced apparent sampling of gate-proximal intermediate states in the mixture simulations than in the C3-only simulations (Fig. 2H, Supplementary Fig. 7F). Together, these results suggest that C7 interferes with C3 access to ECL2-ECL3 gate and subsequent entry into the OBS, meanwhile its own binding leads to nonproductive conformation, supporting the prediction that C7 acts as an antagonist of C3 in the WT OR51E2.

To validate this prediction experimentally, we measured OR51E2 activation in the presence of a range of concentrations of C3 as a function of increasing concentration of C7. OR51E2 activation in the presence of C3 was reduced by C7 with a calculated IC50 of ∼log -1.79, confirming that C7 functions as an antagonist (Fig. 2I, agonist and antagonist tested under various concentrations are shown in Supplementary Figs. 1D,E).

Together, our computational and experimental results support the idea that a ECL2–ECL3 gate-close conformation is required for OR51E2 activity, and C7 serve as an antagonist by keeping the ECL2-ECL3 gate open.

### Allosteric Coupling Between the OBS and ECL2-ECL3 Gate Reverses Odorant Size Selectivity

Extended number of MD simulations (ten runs each 5μs long) of C3 association in F155A and L158A mutant OR51E2 did not result in fully associated state of C3 (Supplementary Table 1). Odorant occupancy density maps show that C3 remains localized near the ECL2–ECL3 gate across all trajectories (Figs. 3A, 3B). Distance measurements of C3 to various residues in the binding site show that although C3 forms contacts with R262, the ECL2–ECL3 gate remains in closed conformation, in contrast to the dynamic “open and close” equilibrium observed in the wild-type simulations (Fig. 3C, green curve; Fig. 3D). Based on these observations, we hypothesize that the F155A and L158A mutations allosterically impair C3 activity by stabilizing a ECL2–ECL3 gate-close conformation.

**Figure 3.**
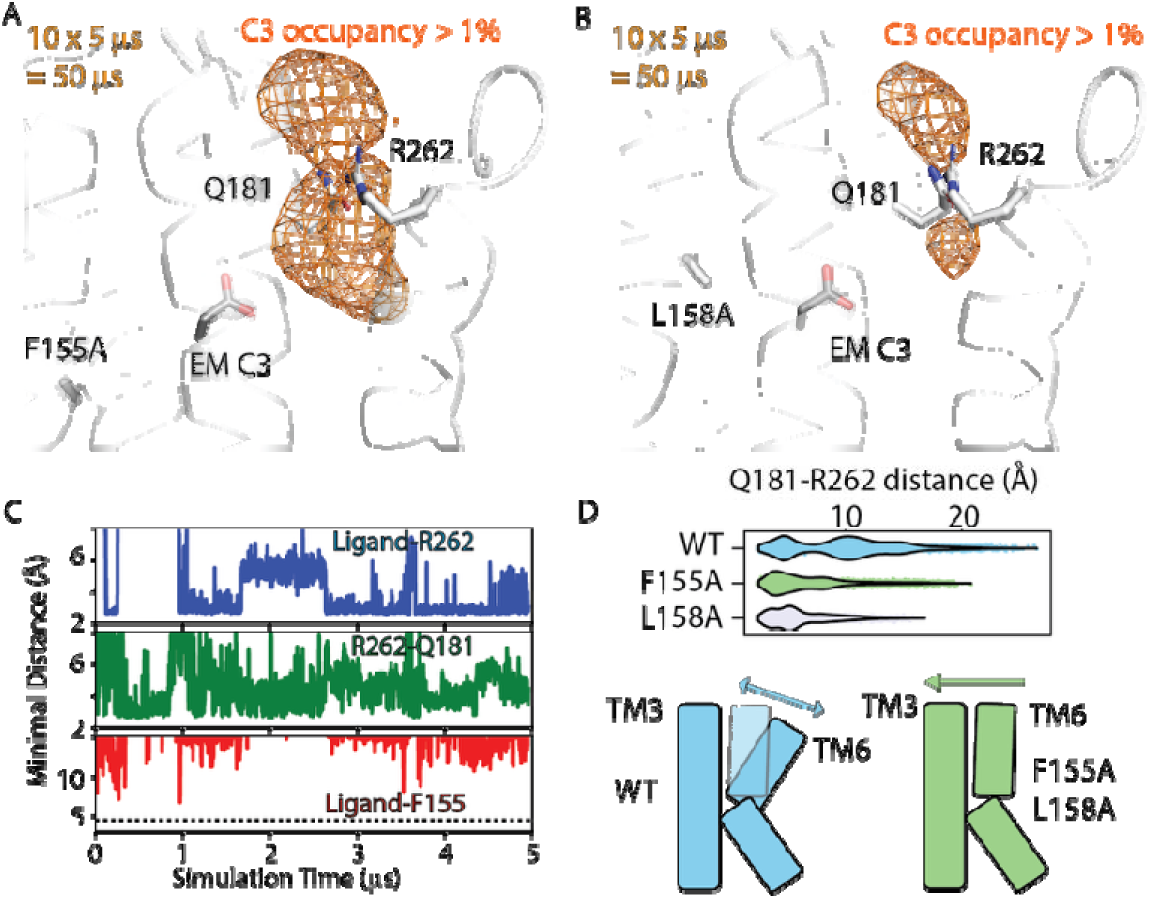
Residues in the bottom of the odorant binding pocket allosterically regulate the opening and closing of the extracellular ECL2-ECL3 gate conferring odorant selectivity. (A,B) Structural regions in OR51E2-F155A and OR51E2-L158A mutants with > 1% C3 occupancy . The binding pose of C3 in the cryo-EM structure is shown as reference. (C) Time series plots of distance of C3 with different residues in the binding site that characterize the odorant association process from C3 association in WT OR51E2 process: Blue: Minimal distance between C3 and R262; Green: Minimal distance between R262 and Q181; Red: Minimal distance between C3 and F155. (D) Violin plot of the minimal distance between Q181 and R262 measured from dissociated state for C3 association into WT, F155A and L158A respectively.

Together, these extended simulations indicate that OBS mutants F155A and L158A allosterically stabilize a closed ECL2–ECL3 gate, associated with persistent C3 sampling near the gate and reduced C3-evoked activity.

### Unlike in class I OR51E2, odorants associate through the membrane interface in OR1A1, a class II OR

Class II ORs represent a major branch of the OR family. One commonly noted distinction between class I and class II ORs, although not universal, is the physicochemical character of their native odorants. Class I ORs often recognize relatively water-soluble carboxylic acid odorants, whereas class II ORs tend to bind a broader and more chemically diverse set of less water-soluble odorants. On this basis, we hypothesized that odorant association might follow different routes in the two classes. To examine this possibility, we performed odorant-association MD simulations for the class II receptor OR1A1 using a structural model derived using our recently published consensus OR1 cryo-EM structure as template.^5^ We focused on two well-studied odorants, L-menthol and R-carvone at their EC50 concentration, and simulated both WT OR1A1 and the G108A variant, which differentially modulates L-menthol and R-carvone evoked activity. Pooling across all four simulation systems with various conditions (L-menthol WT, R-carvone WT, L-menthol G108A, R-carvone G108A, 5 random velocities for each system with 440 μs of simulation in total, see Supplementary Table 2 for details), we observed five distinct odorant-association paths that lead into the OBS of OR1A1 (Figs. 4A, 4B).

**Figure 4.**
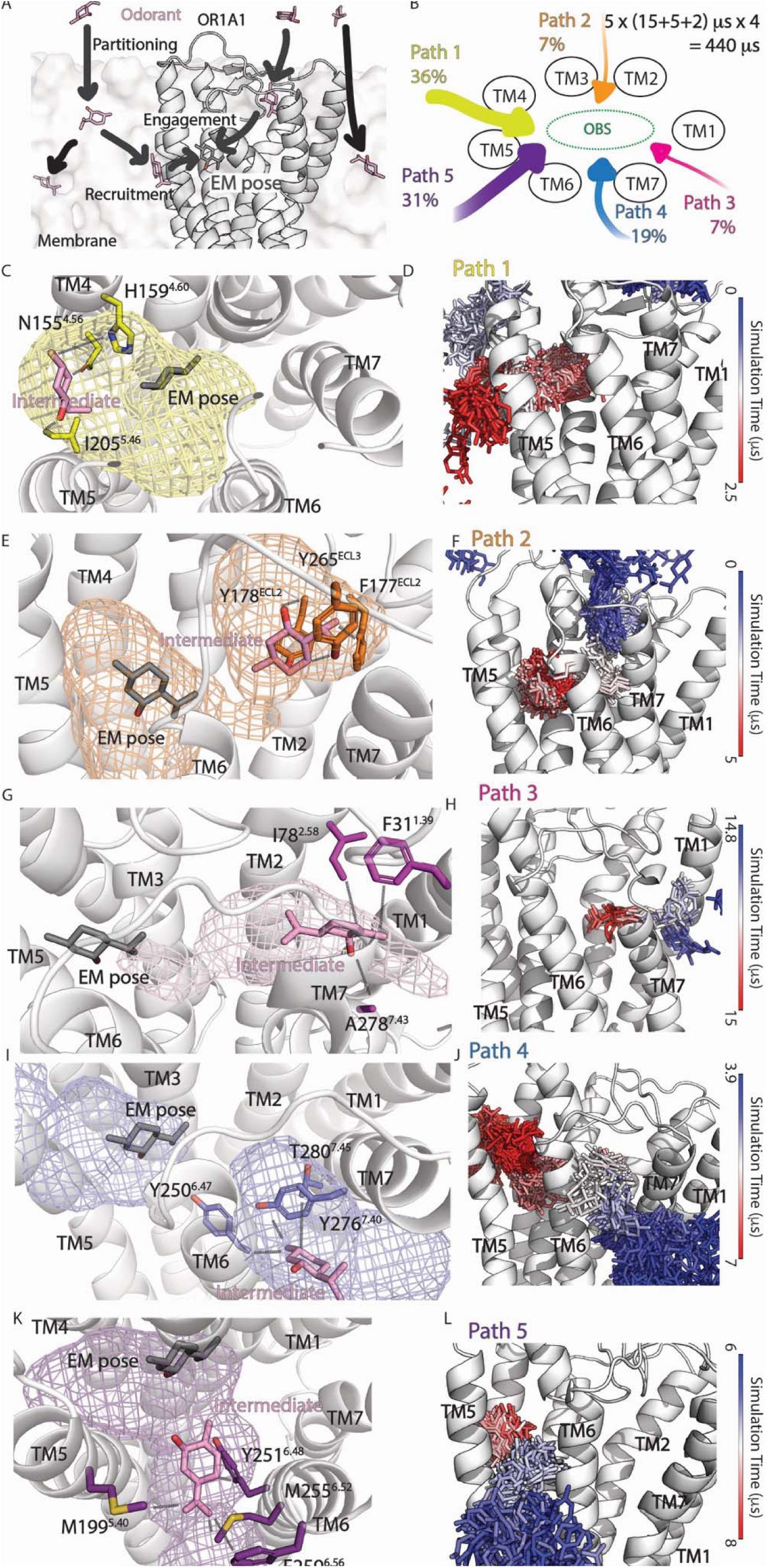
Five odorant association paths and their intermediate gating states in OR1A1. (A) Overview of the multistep odorant association process for OR1A1. Odorants first partition from bulk solvent into the lipid bilayer (partitioning), accumulate near the GPCR–membrane interface (recruitment), and enter the receptor through a gap between TM helices before settling into the odorant binding site (OBS; engagement). In rare cases, odorants access the extracellular vestibule directly from solution via the ECL2–ECL3 gate. The cryo-EM bound pose (“EM pose”, PDB ID: 8UXY) of L-menthol is shown as a reference for the final engaged state. Schematic top-down view of the OR1A1 TM bundle showing the five entry paths identified by pooling association events across all four simulation systems (L-menthol WT, R-carvone WT, L-menthol G108A, R-carvone G108A). Each path is color-coded and labeled with its fractional flux (a probability described by percentage of total observed association events). The OBS is shown as a green dashed ellipse. (C–L) Representative intermediate-state ensembles and time-colored association trajectories for each path, taking one association event out of all events from four simulation box (L-menthol WT, R-carvone WT, L-menthol G108A, R-carvone G108A) as example. Colored mesh volumes depict the spatial occupancy by the odorant during the intermediate state and associated state (>1% of simulation time). The L-menthol EM pose is shown for reference. Key gating residues that contact the odorant at high frequency (>40%) during the intermediate state are labeled with Ballesteros–Weinstein numbering. (D, F, H, J, L) Time-colored association trajectories of corresponding association events. Individual odorant conformations sampled every 20 ns are rendered as sticks, colored from red (early in trajectory) to blue (late), showing the progression of the odorant from the membrane or extracellular space through the gate and into the OBS. Time scale of chosen association example are labeled in color bar.

The five paths differ in their entry geometry relative to the TM helices and probability of utilizing the path (Fig. 4B). Path 2 with a probability of 7%, involves entry from the extracellular space through an ECL2–ECL3 gate, resembling the extracellular paths observed for OR51E2. In the remaining four pathways, odorants first partition from bulk solvent into the lipid bilayer and subsequently access the OBS through gaps between TM helices: path 1 via the TM4–TM5 interface with a probability of 36%, path 3 via the TM1–TM7 interface with a probability of 7%, path 4 via the TM6–TM7 interface with a probability of 19%, and path 5 via the TM5–TM6 interface with a probability of 31% (One example out of all events is shown in Figs. 4C–L; More details in Supplementary Table 2). The predominance of membrane-mediated entry in OR1A1 is broadly consistent with the hydrophobic character of its odorants, though this interpretation does not exclude other contributing factors.

Each path is associated with a distinct intermediate state in which the odorant dwells at the corresponding TM gate prior to full engagement with the OBS. The characteristic gating residues (see method for details) framing each intermediate were identified as those with odorant contact frequencies greater than 40% within the intermediate ensemble. For path 1, the gate is framed by residues H159^4.60^, N155^4.56^, and I205^5.46^, which form the TM4–TM5 portal (Fig. 4C, 4D). Path 2 intermediates localize near Y265^ECL3^, Y178^ECL2^ and F177^ECL2^ at the extracellular vestibule (Fig. 4E, 4F). Path 3 passes through residues in the TM1–TM7 crevice, with odorant contacts dominated by F31^1.39^, I78^2.58^ and A278^7.43^ (Fig. 4G, 4H). Path 4 intermediates engage Y250^6.47^, Y276^7.40^ and T280^7.45^ at the TM6–TM7 interface (Fig. 4I, 4J), while path 5 intermediates are coordinated by M199^5.40^, M255^6.52^, Y251^6.48^, and F259^6.56^ at the TM5–TM6 gate (Fig. 4K, 4L). We noticed few gate residues are classified as OBS residues in previous work,^5^ we acknowledge such overlap to different categorize method. We mutated every gating residue to Alanine (or Leucine when it is Alanine in WT) and measured the activity of both R-carvone and L-menthol using heterologous cell-based assay and showed that majority of the gating residues indeed lead to change in odorant mediated responses (Supplementary Fig. 3).

Taken together, both simulation and mutagenesis results showed that OR1A1 supports at least five geometrically distinct entry routes, four of which are membrane-mediated, and each is gated by a characteristic set of TM helix residues. This stands in contrast to OR51E2, where odorant association was observed through extracellular paths. The diversity of membrane-accessible gates in OR1A1 likely reflects the hydrophobic nature of its odorants, which preferentially partition into the bilayer and thereby gain access to a larger receptor surface area compared with water soluble odorants in OR51E2.

### Gating residues tune OR1A1 odorant specificity and are allosterically biased by G108A mutant

Having established the five association paths and their gating residues, we next asked whether these paths are used differently by L-menthol and R-carvone, which are related to odorant specificity. We first compared the total number of observed odorant association events per path for L-menthol and R-carvone in the WT receptor (Figs. 5A, 5B, including both regular and enriched MD, 5 random velocities, a total of 110 μs trajectories for each of the two odorants). Strikingly, R-carvone showed more observed association events than L-menthol across the simulation ensemble, suggesting a higher apparent association propensity under the sampled conditions. MSM analysis was consistent with this qualitative trend by identifying recurrent intermediate and associated states occupancies (Supplementary Fig. 4A, 4B). Thus, the difference in overall association propensity represents one notable distinction between the two odorants in the WT receptor.

**Figure 5.**
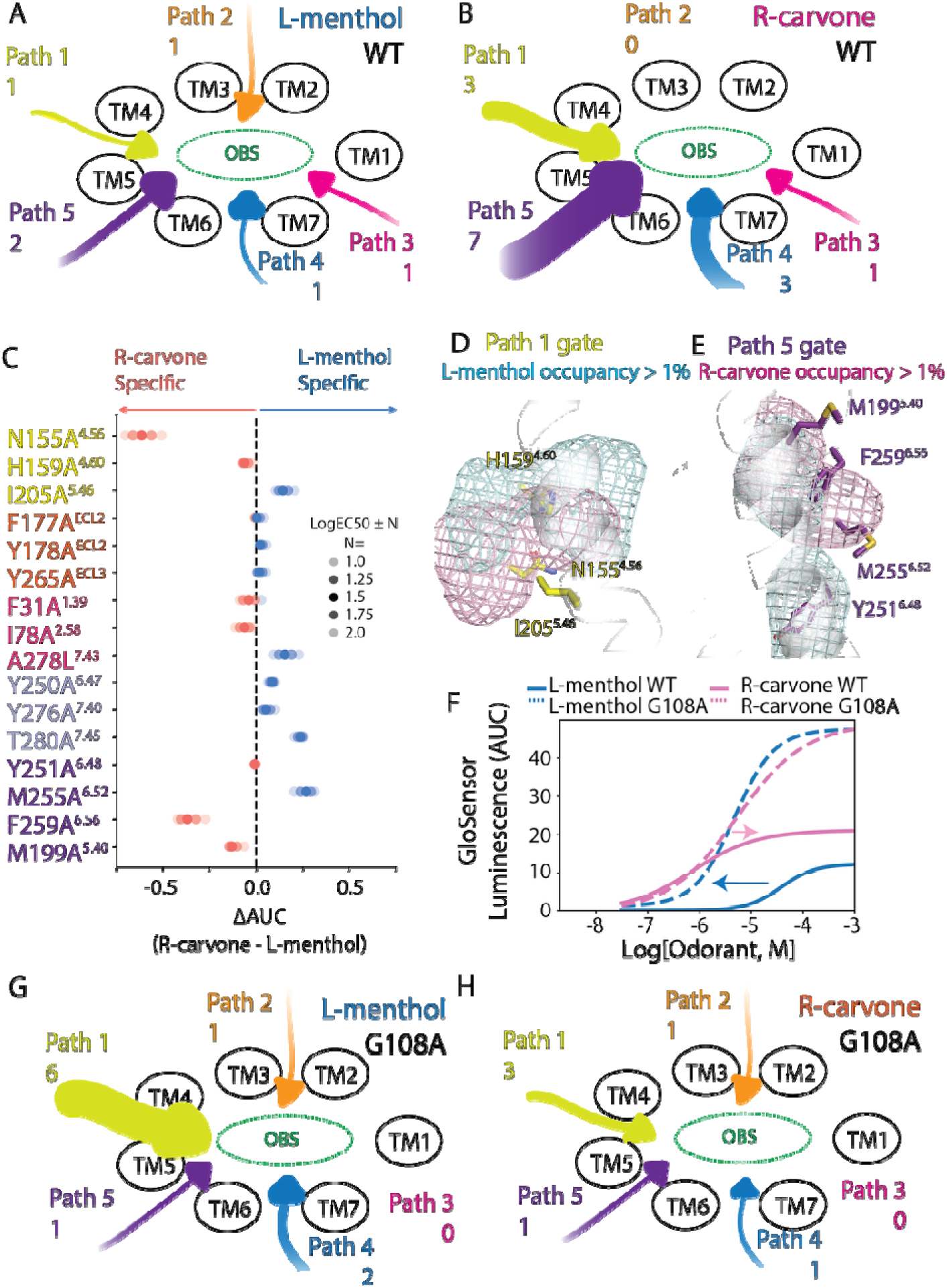
Odorant-specific path utilization, gating residue contributions, and allosteric effects of G108A in OR1A1. (A, B) Schematic diagrams showing the number of observed odorant association events per path for (A) L-menthol and (B) R-carvone in OR1A1 WT. Arrow size and the number adjacent to each path label indicate the count of association events. (C) Bubble plot of gating residue mutant effects on odorant specificity. The x-axis shows ΔAUC (R-carvone – L-menthol) from GloSensor cAMP assays following alanine substitution (or leucine when the WT residue is alanine) of each gating residue, with implied odorant selectivity labeled. Dot transparency encodes the ΔAUC estimated under different concentration windows. (D, E) Structural view of the path 1 (D) and path 5 (E) gate intermediate state. The pink mesh shows the R-carvone-occupied volume (>1% MD frames) during the intermediate state from all observed R-carvone association events in WT, while the blue mesh is the occupancy volume of L-menthol from all observed L-menthol association events in WT. (F) Experimental dose–response curves for L-menthol (blue) and R-carvone (pink) in OR1A1 WT (solid lines) and G108A mutant (dashed lines), measured by GloSensor luminescence. Arrows show the potency change from WT to G108A. (G, H) Schematic diagrams showing per-path association event counts in OR1A1 G108A for (G) L-menthol and (H) R-carvone.

To test whether gating mechanisms contribute to odorant discrimination, we quantified odorant specificity from gating residue mutagenesis results (Supplementary Fig. 3). We calculated the resulting change in odorant-evoked responses as ΔAUC (difference in the area under the dose dependent curves; L-menthol − R-carvone; Fig. 5C). Across the 16 residues tested, several mutations produced significant and directional shifts in ΔAUC. N155A^4.56^, a gating residue at the path 1 TM4–TM5 gate, yielded a negative ΔAUC, indicating a larger change in R-carvone activity and identifying N155 as an R-carvone-specific gating residue. Similarly, F259A^6.56^ behaved as a path 5 R-carvone-specific gating residue, whereas M255A^6.52^ within the same path 5 gate produced an L-menthol-specific effect. Notably, these odorant-specific gating residues cluster in paths 1 and 5, consistent with our MD results showing that paths 1 and 5 are more frequently used for odorant association (Fig. 4B).

We next asked why N155^4.56^, M255^6.52^, and F259^6.56^ show odorant-specific effects in the gating residue mutational analysis. We pooled all observed odorant association trajectories for paths 1 and 5 for both L-menthol and R-carvone, and we computed odorant occupancy over intermediate state frames. This analysis revealed distinct intermediate-state spatial distributions for the two odorants. In path 1 intermediates, R-carvone density was enriched and localized around N155^4.56^ (Fig. 5D). In path 5 intermediates, R-carvone similarly occupied the TM5–TM6 gate region near F259^6.56^, whereas L-menthol density was shifted toward M255^6.52^ (Fig. 5E). Together, these odorant-specific intermediate occupancies are consistent with a model in which gating residues tune selectivity by stabilizing different ligand-dependent intermediate ensembles and thereby modulating gate residence before productive engagement of the orthosteric site.

T280^7.45^, which lies on path 4, appears more L-menthol–specific than R-carvone (Fig. 5C). However, this mutation produces only a slight change in L-menthol potency and has little effect on R-carvone (Supplementary Fig. 3). This small shift makes it unclear whether the resulting ΔAUC difference reflects a meaningful, biologically relevant selectivity effect. Y251^6.48^ on path 5 sits near the L-menthol intermediate occupancy volume which make it seems like a L-menthol specific residue (Fig. 5E), but Y251A^6.48^ eliminates signaling for both odorants to a similar extent. The lack of odorant specificity in ΔAUC is therefore most consistent with Y251^6.48^ serving a general activation role in OR1A1 rather than acting as a determinant that preferentially supports L-menthol.

We next introduced the G108A mutation, a substitution within the OBS that is known to increase L-menthol potency while decreasing R-carvone potency in cell-based assays (Fig. 5F). To understand whether odorant association might be affected by this OBS mutation, we count association event per path in G108A for each odorant (Figs. 5G, 5H, including both regular and enriched MD, 5 random velocities, a total of 110 μs trajectories for each system). G108A markedly increased the number of L-menthol association events observed across the simulation ensemble, while simultaneously reducing R-carvone association events. This directional shift in association frequency mirrors the experimental potency changes: L-menthol becomes more potent and R-carvone less potent in G108A. These simulation observations are consistent with the MSM-derived apparent state distributions, which suggest opposite effects of G108A on L-menthol and R-carvone association: enrichment of associated-like L-menthol states and reduced associated-like sampling for R-carvone (Supplementary Fig. S4C, S4D).

Together, the agreement between MD-observed association frequency and experimental AUC data suggests that allosteric effects of G108A on gate dynamics may contribute, at least in part, to the experimentally observed changes in odorant potency. We emphasize that odorant association represents one component of the overall receptor activation process, and that changes in association frequency alone do not fully account for all aspects of potency or efficacy. Nonetheless, the consistent directional agreement between simulation and experiment supports the idea that OBS mutations can allosterically rewire the entry gates to differentially favor one odorant over another, providing a plausible mechanism by which orthosteric substitutions reshape odorant specificity in OR1A1.

### Like OR51E2, an OR1A1 antagonist occupies gate region more often than the agonist

In OR51E2, our C3–C7 mixture simulations showed that antagonist C7 has higher intermediate state population than agonist C3. We asked whether this behavior extends to OR1A1. Using an established OR odorant library with diverse physicochemical properties, we identified (+)-limonene as an agonist and 3,4-hexanedione as an antagonist of OR1A1 (Fig. 6A, 6B, Supplementary Fig. 5). We then performed multi-odorant mixture MD simulations and observed a similar pattern as in OR51E2. Odorant occupancy maps revealed pronounced density at gates, with 3,4-hexanedione forming occupancy clouds at the entrances of all five paths, whereas (+)-limonene showed weaker localization that was largely restricted to the gates of paths 1 and 2 (Fig. 6C). Consistent with these observations, MSM analysis indicated greater enrichment of the antagonist in intermediate states (Fig. 6D, 20% for 3,4-hexanedione versus 8% for (+)-limonene). On the simulated timescales, no agonist association events were observed, whereas the antagonist reached the associated states with a population of 21%, potentially due to higher occupancy at the gate.

**Figure 6.**
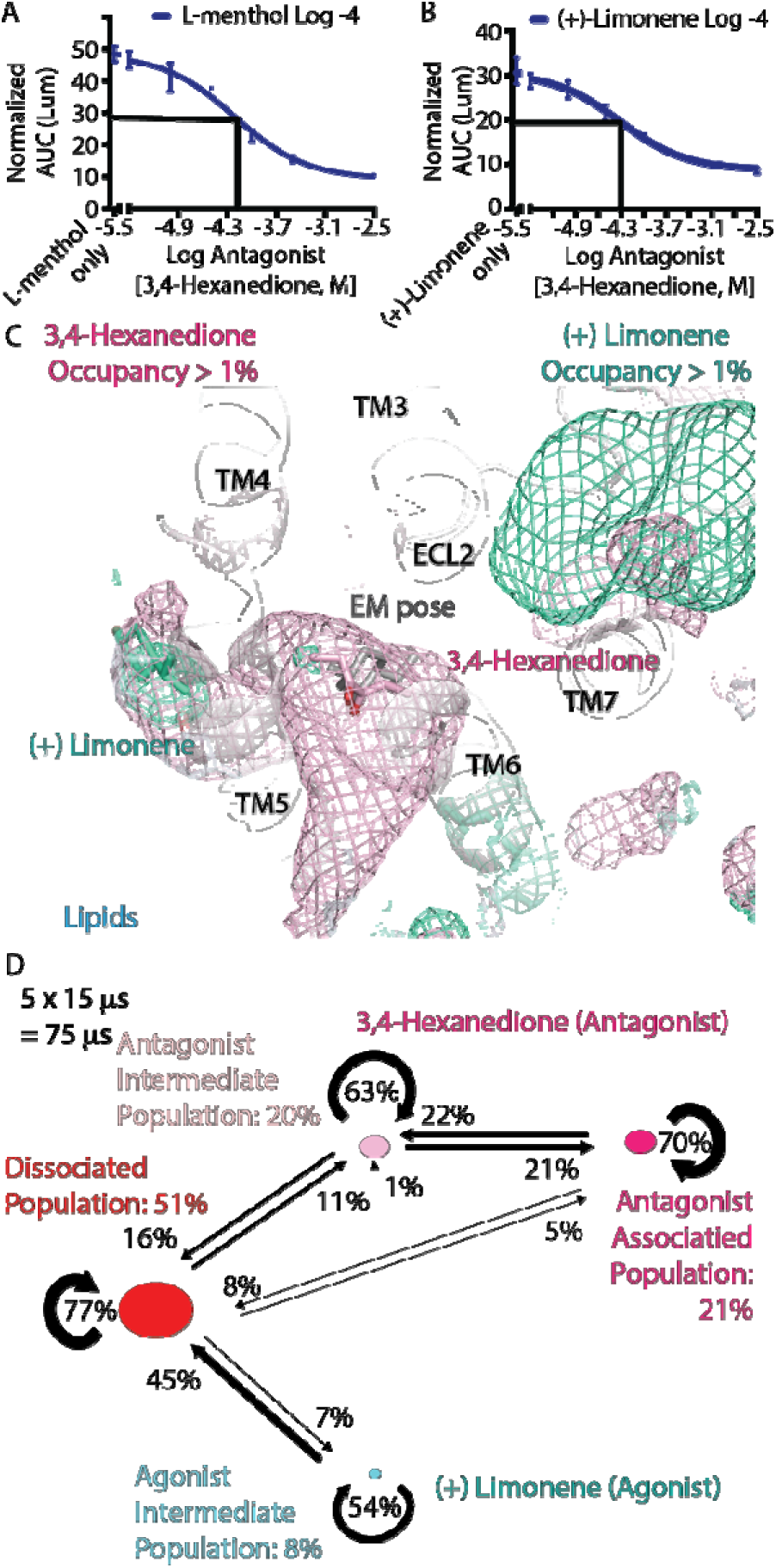
The antagonist 3,4-hexanedione occupies gate more often than agonist (+)-limonene. (A) Antagonist titration experiment with L-menthol (agonist) fixed at 10^−4^ M. The plot shows normalized GloSensor AUC versus 3,4-hexanedione concentration. (B) Antagonist titration experiment with (+)-limonene (agonist) fixed at 10^−4^ M. The plot shows normalized GloSensor AUC versus 3,4-hexanedione concentration. (C) Odorant occupancy around OR1A1 during mixture simulations. 3,4-hexanedione and (+)-limonene are shown as occupancy meshes at a 1% threshold. The receptor is shown as a gray cartoon with key structural elements labeled. The cryo-EM L-menthol pose is shown as gray sticks for reference. Lipids are shown for context. (D) Markov state model summary for the mixture simulations. The network reports state populations and transition probabilities. States include dissociated, agonist intermediate, antagonist intermediate, and antagonist associated. An agonist associated state is not observed in these simulations.

Together, our new experiments and simulations support the gating mechanism established in the OR51E2 study, in which antagonists occupy the gate region more often than agonist.

## Discussion

Odorant selectivity is often explained using binding pocket complementarity (the “lock and key” model). Recent odorant receptor structures show why that view is incomplete. These structures also highlight the importance of extracellular loops and access control.^4,20^ Our MD simulation results and mutagenesis data support a mechanistic model in which odorant entry and intermediate-state residence may contribute to selectivity in both a class I receptor OR51E2 and a class II receptor OR1A1.

Our OR51E2 simulations extend prior structural and functional work by defining a mechanistic role for the ECL2-ECL3 gate in odorant access. Previous studies established that extracellular loop regions are critical for shaping the orthosteric pocket and regulating odorant responsiveness, but the microscopic steps by which these loops control ligand entry remained unresolved. Here, the simulations suggest that C3 and C7 approach OR51E2 through extracellular routes that converge near the ECL2–ECL3 gate, where gate closure is associated with productive odorant transfer into the orthosteric pocket. This gate-centric model is consistent with prior work. ^4,21^ Importantly, our results further suggest that this gate can be allosterically biased by odorant identity and receptor mutations, providing a microscopic entry-gate mechanism for antagonism.^22^

For OR1A1, our simulations support a different access architecture. Odorants partition into the lipid bilayer and reach the binding site through multiple transmembrane entry paths. Prior GPCR studies and simulations support membrane-mediated entry for hydrophobic ligands in multiple other receptor systems.^23^ A related example shows that a GPCR can support both a membrane-facing channel and an extracellular-facing channel for ligand entry.^24^ A recent insect odorant receptor study also reports dual solvent-facing and membrane-facing paths in a chemosensory receptor.^25^ Insect odorant receptors are ion channels, so the architecture differs. The pathway concept is still useful because it shows how chemosensory receptors can support more than one access route.

Our OR1A1 results also connect to the OR1A1 mutagenesis literature. Prior work identifies Gly108 and Asn109 on TM3 as key determinants for carvone enantiomer selectivity and binding geometry.^5,26^ Recent OR1A1 structure-activity work that combines ligand series, docking, and molecular dynamics reinforces the importance of these determinants and summarizes earlier mutagenesis results.^27^ Our data supports a model linking between an orthosteric substitution and odorant association path control. It supports an allosteric model in which the orthosteric site biases gate and pathway ensembles.

Finally, our OR1A1 mixture simulation results suggest that the antagonism principle can generalize across receptor classes. The antagonist 3,4-hexanedione occupies gate regions more often than the agonist (+)-limonene under same concentration. This parallels the OR51E2 result. It supports a shared mechanism that depends on intermediate occupancy. It also suggests a possible design principle for antagonist development. Molecules that increase gate residence while disfavoring productive engagement are plausible antagonists.

This study has limitations. We quantify association and intermediate residence, but we do not simulate the full activation trajectory through G protein coupling. We also rely on finite sampling and elevated ligand concentrations to capture spontaneous events. These constraints limit direct interpretation of absolute state populations as thermodynamic observables. Future work can test the gating mechanism using loop distance reporters, such as FRET/BRET, targeted gate mutations that tune closure probability, and lipid composition perturbations that bias membrane-facing portals.

In summary, entryl1lpath choice and gatel1lresidence occupancy may contribute to agonism, antagonism, and odorant selectivity across ORs under study. By shifting attention from static pockets to dynamic access, this work lays a foundation for gatel1ltargeted antagonists, positive allosteric modulators that bias entrance, and environmental strategies (such as membrane tuning) to engineer receptor responses in both basic and applied olfaction.

## Methods

### System preparation and MD protocols

All systems were built in CHARMM-GUI^28–30^ under the odorant concentrations and lipid compositions described in Supplementary Table 1. All-atom MD simulations were run with GROMACS 2021^31^ using the CHARMM36m force field.^32^ Odorant parameters were generated with the CHARMM General Force Field (CGenFF),^33^ and lipid parameters were taken from the CHARMM36m lipid library. Receptor protonation states were assigned with the Maestro Protein Preparation Wizard with pH set to 7.0. Systems were solvated with TIP3P water and neutralized with 0.15 M KCl. Final box dimensions were ∼80 × 80 × 135 Å for OR51E2 and ∼100 × 100 × 110 Å for OR1A1. The initial conformation for simulation of OR51E2 (PDB ID: 8F76) are taken from previous study, with homology modeling method, loop refinement, sidechain refinement method described.^4^ As for the initial conformation for simulation of OR1A1, we construct OR1A1 homology model by Swiss-Model,^34^ using the consensus OR1 structure template (PDB ID: 8UXY).

After energy minimization with harmonic positional restraints (10 kcal mol⁻¹ Å⁻²) on heavy atoms of receptor, odorant, and lipids, systems were heated from 0 K to 310 K over 1 ns in the NVT ensemble (Nosé–Hoover thermostat),^35^ then equilibrated for 1 µs in the NPT ensemble (Parrinello–Rahman barostat at 1 bar). During heating and the initial phase of equilibration, restraints of 10 kcal mol⁻¹ Å⁻² were applied for 1 ns and then reduced to 5/4/3/2/1/0 kcal mol⁻¹ Å⁻² in 5-ns increments, after which all atoms were unrestrained for the remainder of the equilibration. Production MD comprised five independent replicas per system to simulation time lengths detailed in Supplementary Table 1, starting from the final equilibrated snapshot, each with distinct random seeds.

A 2-fs time step was used. Bonds to hydrogens were constrained with LINCS (water molecules constrained with SETTLE). Short-range nonbonded interactions employed a 12 Å cutoff, long-range electrostatics were treated with particle-mesh Ewald (PME).^36^ Trajectories were saved every 20 ps. Visualization used VMD and PyMOL (v2.0, Schrödinger). Analyses were performed with MDAnalysis and PyEMMA; contact frequencies were computed with Get_Contacts, and structural figures were generated in PyMOL. A detailed descriptions of all simulations in this study is provided in Supplementary Table 1.

### MSM construction

We built a MSM for each system in PyEMMA using the following workflow.

1. **Feature definition (odorant-residue distance array).** Whole-trajectory odorant–protein contact frequencies were first computed with Get_Contacts. GPCR residues with >20% contact frequency with any odorant were designated “contacting residues.” Using MDAnalysis, we built a Universe from the system PDB and aligned trajectory files, selected heavy atoms of odorants and the contacting residues and compute minimum distances between each odorant and each contacting residue. Distances < 4.5 Å were recorded, values ≥4.5 Å were set to a large sentinel 999 to denote “no contact”. The resulting data were organized into a 3D array (residues × odorants × frames), flattened to a 2D matrix (features × frames), and used as MSM input (Supplementary Fig. 6A). The contacting residues in OR51E2 include R262, H104, F155, I202, S258, Q181, H261, A201, M206, R272, L254 and A108 (Supplementary Fig. 6A left panel). The contacting residues in OR1A1 include M104, I105, G108, N109, D111, S112, N155, H159, I181, M199, I205, F206, L201, L162, L185, F194, M199, V203, Y250, Y251, F259, Y258, T253, V254, Y276, T277, Y178, F177, A278, T280, N176, Y265, V17, F31, H85, F27, T77, M81, I78, F73 and S74.
2. **Hyperparameter scan.** We performed a hyperparameter scan over lag times (τ ∈ {25–500} frames) and numbers of clusters (K ∈ {2–19}) to select an optimized (τ, K) per system. Using the 2D matrix from step 1 as a single trajectory, we ran TICA at each τ (dim=2), clustered the 2D projections with k-means (K=10) and estimated a reversible MSM at the same τ; implied timescales ti (r) = -r/ ln Ui were computed, padded with NaNs for alignment, and plotted versus τ on a semilog y-axis, where plateaus of the slowest curves indicate approximate Markovianity. To assess discretization sensitivity, we fixed τ=300, featurized by TICA, clustered with k-means for K=2 to 19 (fixed seed), built MSMs, and plotted ti (K) versus K (one curve per rank-ordered timescale). We chose K where leading timescales and inferred metastable dimensionality stabilized with increasing K, noting that rank-matching across K does not guarantee mode identity. Across systems, τ≈250–300 and K≈10 or 14 were typically favored (Supplementary Fig. 6B, 6C).
3. **Macrostate definition.** Using the selected (τ, K), we discretized trajectories and built MSM microstate transition network over K microstates (Supplementary Fig. 6D right panel). Centroid structures for each microstate were saved. Overlaying all centroids revealed 2 different odorant-association pathways for C3 and C7 into OR51E2 (C3: Supplementary Fig. 6D left panel, C7: Supplementary Fig. 6G), and 5 different pathways for L-menthol and R-carvone into OR1A1 (Supplementary Fig. 6H). To illustrate the real association process rather than aggregate statistic, we extracted the trajectory that captured the event and generated time-series visualizations (Fig. 1A, 2A, 2B, 4D, 4F, 4H, 4J, 4L, Supplementary Fig. 2A-E).
4. **Microstate assignment.** We coarse-grained the microstate MSM into a small set of metastable macrostates using PCCA+,^37^ then applied minimal expert curation to align labels with pathway biology. Consequently, OR51E2 was represented with four macrostates (Associated, two pathway-specific Intermediates, Dissociated) to distinguish its two association paths, whereas OR1A1 was summarized with three macrostates (Associated, Intermediates, Dissociated) to emphasize global behavior and enable cross-odorant comparison. Implementation and visualization used PyEMMA, following standard MSM coarse-graining practice in biomolecular kinetics.^38^
5. **Reduced MSMs and transitions.** From the PCCA+ guided mapping, we built reduced MSM and reported macrostate transition probabilities together with macrostate populations as the primary network summary for comparing odorant occupancies across odorants/conditions with an established MSM protocol in PyEMMA (Fig. 1C, 2C, 2H, 6D, Supplementary Fig. 4, 7F).^39^

### Odorant Occupancy map calculation

Odorant occupancy maps were computed with VMD’s VolMap tool. Trajectories were imaged under PBC, rewrapped, and recentered on the receptor so that OR coordinates remained fixed across frames. Using the full trajectories, we calculated occupancy of odorant heavy atoms. We calculated the occupancy odorant heavy atoms after alignment by OR backbone, using VMD volmap tool. The occupancy map was generated over all frames on a 1.0 Å grid over the average. Each voxel of the grid records the fraction of frames (1% cutoff was used in this study) in which at least one odorant heavy atom occupies said voxel. Isosurfaces were rendered in PyMOL as mesh objects with isovalue ≥ 1% over the simulation (Fig. 1E, 2F, 3A, 3B, 4C, 4E, 4G, 4I, 4K, 5D, 5E, 6C).

### Distance measurements

We computed distances from MD trajectories with MDAnalysis. Distances were calculated in angstrom units using Euclidean geometry. For each frame, we computed an all to all distance matrix between two atom selections (only non-Hydrogen atoms were selected) and then reported the minimum value as the distance for that frame. This minimum distance definition was used both for single distances shown as time series and for the odorant residue distance arrays used in MSM construction.

### Gate distance and odorant association distances

To monitor the ECL2–ECL3 gate in OR51E2, we measured the Q181–R262 distance as the minimum distance between Q181 side-chain atoms OE1 and NE2 and R262 side-chain atoms NH1 and NH2. For odorant capture and transfer steps, we measured the minimum distance between the odorant head group atoms and the R262 NH1 NH2 atoms. We measured the progress of full association into the orthosteric site by the minimum distance between odorant heavy atoms and selected pocket reference residues such as F155. These distances were plotted as time series using the frame index and the effective time spacing of the saved trajectory frames (Fig. 1F, 1G, 2D, 2E, 2G, 3C, 3D, 4C, 4E, 4G, 4I, 4K, Supplementary Fig. 2F-I).

### Odorant docking

We docked C6, C7, and C8 into the OR51E2 L158A and F155A mutants. Alanine substitutions were introduced in Maestro. Docking grids were generated with Glide Receptor Grid Generation, centered on the cryo-EM C3 pose, with a 20 Å margin box. Carboxylic acids were prepared in Maestro’s odorant preparation workflow to assign charges and rotatable bonds. Docking was performed with Glide in extra-precision (XP) mode using default settings. For each odorant, the top-scoring pose was retained as the reference for Supplementary Fig. 2F-I.

### cAMP signaling assays Cells and transfection

GloSensor cAMP assays (Promega) were employed to measure the cAMP levels in HEK293T cells downstream of odorant receptor activation, as previously described.^40^ HEK293T or Hana 3A cells are cultured in minimum essential medium (MEM; Corning) supplemented with 10% FBS (Gibco), 0.5% penicillin-streptomycin (Gibco) and 0.5% amphotericin B (Gibco). The cells were plated the day before transfection at 1(8 × 10 –1×10) cells/ml. For each 96-well plate, 1,000 ng pGlosensor-20F plasmid (Promega), 500 ng RTP1S plasmid, and 7,500 ng of rho-tagged OR in the pCI mammalian expression vector (Promega) were transfected with 20µL of Lipofectamine 2000 (Invitrogen) in MEM supplemented with 10% FBS. After 18-24 h incubation, the transfection media were replaced with 25 µL of 2.6% GloSensor substrate (Promega) solution diluted in HEPES buffer supplemented with D-glucose. The plates were incubated at room temperature for 2h.

### Odorant preparation

For the entry path experiments, the odorant stocks (0.1M or 1M in ethanol) were diluted on the day of the assay into HBSS containing D-glucose to reach the indicated final concentrations. For antagonism competition assays, stock solutions of both agonists and antagonists were freshly diluted the day of the experiment, to create a dose response of dose response for the agonist challenged with antagonist, as follows. A starting saturation stock concentration of both the agonist and antagonist is prepared, separately. Half-log dilutions from the starting concentrations are made in HBSS containing D-glucose to make 5 agonist concentrations and 7 antagonist ones. The antagonist and agonist are added to the cells sequentially with no antagonist pre-treatment.

### Luminescence acquisition and AUC quantification

Luminescence was measured for 15 min immediately after odor stimulation using the CLARIOstar^Plus^ plate reader (BMG LABTECH). We computed cAMP accumulation by summing luminescence values across the measurement cycles. For the entry path experiments, the cAMP accumulation value was normalized by the empty vector control and basal activity was subtracted. Normalized responses were defined as the ratio to the WT maximum. Dose–response curves were fitted using four-parameter logistic regression in Prism 10.6.1 (GraphPad Software). The area under the dose-response curve (AUC) for each receptor was calculated from the fitted curve over the range of logEC50 ± 1.5 log unit of the WT control. The relative AUC was defined as AUC_mutant_ / AUC_WT_. Data represent 3-5 independent experiments with three technical replicates per condition. Error bars indicate standard deviation (SD).

### Antagonism and mixture assays

For antagonism competition assays, both odorants (agonist and antagonist) were added sequentially within seconds of each other. Using multiple fixed agonist concentrations and antagonist concentrations titrated against to produce a single inhibition dose response curve for each agonist concentration. cAMP activity using the Glo sensor assay, as previously described, following odor stimulation was normalized to basal activity (cAMP accumulation in the absence of odorant) for each concentration. The response at each concentration was calculated as the sum of the luminescence signal across all recorded measurement cycles, producing an integrated response value to create an AUC data point for each concentration mixture. Dose–response curves were fitted using three-parameter logistic regression in Prism 10.4.2 (GraphPad Software), from which log EC50 (for agonist only) and log IC50 (for antagonist at ∼ EC50 agonist concentration) values were obtained. The data represent 3-5 independent experiments with two technical replicates per condition. Error bars indicate the standard error of mean (SEM).

### Mutant preparation

Point mutations were introduced by PCR-based site-directed mutagenesis. For F155A and L158A in OR51E2 and N155A, H159A, Y178A, M199A, I205A, Y251A, M255A, F259A, Y276A, A278L, and T280A in OR1A1, mutagenesis was performed as previously described.^41^ Briefly, mutant constructs were generated by two-step PCR using Phusion polymerase (Thermo Fisher Scientific). The first round of PCR generated two fragments, one containing the 5′ region upstream of the mutation site and the other containing the 3′ downstream region. The second PCR amplification joined these two fragments to produce a full open reading frame of the OR. PCR products of the expected size were gel-purified and cloned into the MluI and NotI sites of the pCI vector that contains rho-tag. For F31A, I78A, F177A, Y250A, and Y265A in OR1A1, PCR fragments were assembled using NEBuilder® HiFi DNA Assembly kits (New England Biolabs). Plasmids were purified using a ZymoPure miniprep kit (Zymo Research). All constructs were verified by Sanger sequencing. Mutants tested in this study included OR51E2 (F155A, L158A) and OR1A1 (F31A, I78A, N155A, H159A, F177A, Y178A, M199A, I205A, Y250A, Y251A, M255A, F259A, Y265A, Y276A, A278L, and T280A).

### OR expression

Flow cytometry was used to evaluate cell surface expression of ORs as previously described.^42^ HEK293T cells were seeded onto 35-mm plates (Corning) with approximately 2.5% confluency. The cells were cultured overnight. After 18–24 h, 1,000 ng of ORs tagged with the first 20 amino acids of human rhodopsin (rho-tag) at the N-terminal in pCI mammalian expression vector (Promega), 200 ng of RTP1S and 10 ng eGFP were transfected using Lipofectamine 2000 (11668019, Invitrogen). At 18–24 h after transfection, the cells were detached and resuspended using Cell Stripper (Corning) and then transferred into 5-ml round-bottom polystyrene tubes (Falcon) on ice. The cells were centrifuged at 4 °C and resuspended in PBS (Gibco) containing 15 mM NaN3 (Sigma-Aldrich) and 2% FBS (Gibco). The cells were stained with 1/400 (v/v) of primary antibody mouse anti-rhodopsin clone 4D2 (MABN15, Sigma-Aldrich) and incubated for 30 min, then washed with PBS containing 15 mM NaN3 and 2% FBS. The cells were stained with 1/200 (v/v) of the phycoerythrin-conjugated donkey anti-mouse F(ab′)2 fragment antibody (715-116-150, Jackson Immunologicals) and incubated for 30 min in the dark. To label dead cells, 1/500 (v/v) of 7-amino-actinomycin D (129935, Calbiochem) was added. The cells were analyzed using a BD FACSCanto II flow cytometer. Gating was performed to select single, spherical, viable and GFP-positive cells. Phycoerythrin fluorescence intensities were analyzed and visualized using FlowJo (v.10.8.1). Results of expression is shown in Supplementary Fig. 9.

## Data Availability

MD simulation trajectories associated with this manuscript is available through Zenodo with DOI: 10.5281/zenodo.19363176 .

## Acknowledgements

We thank Dr. Sneha Bheemireddy for generating an initial draft of the Introductions section of the manuscript. This work was supported by the National Institutes of Health (NIH) grant R01DC020353 to H.M., N.V. and A.M., R35-GM156498 to N.V., and K01DC022958 to N.M.. The computational resources were provided by COH/TGen High Performance Computing cluster (HPC), including Gemini and Apollo.

## Author Contributions

N.M., M.M., N.V., and H.M. designed the study. N.M. performed the MD simulations and analysis. M.M. and D.T. performed the mutation and functional assay experiments. N.M., M.M., D.T., C.L.D.T and J.A. did the analysis of computational predictions and experimental results analysis. N.M., M.M., and N.V wrote an initial draft of the manuscript and generated figures with contributions from all authors. Further edits to the manuscript were provided by A.M. and H.M. The overall project was supervised and funded by N.V., H.M., and A.M.

