## Supplementary Material for "Allosteric Gating Mechanism Regulates Odorant Selectivity and Antagonism in Odorant Receptors"

**Supplementary Information**

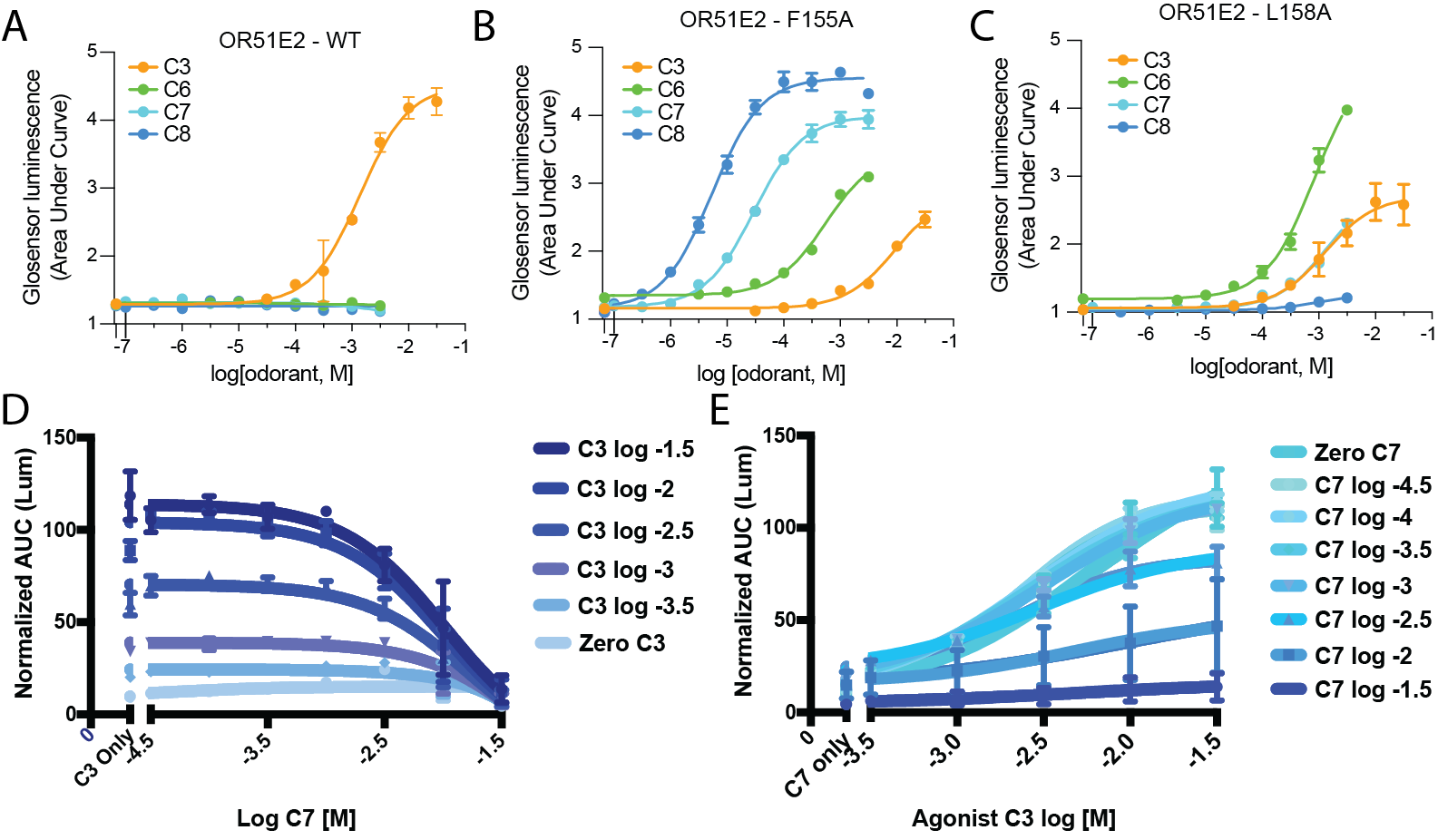

**Supplementary Figure 1. Odorant-evoked OR51E2 activation.** The area under the GloSensor luminescence trace (AUC) quantifies receptor activation by propionate (C3), hexanoate (C6), heptanoate (C7), and octanoate (C8) in (A) OR51E2 WT, (B) OR51E2-F155A, and (C) OR51E2-L158A. (Panel A-C are adapted from reference 4) (D) C7 titration at various fixed C3 concentrations in OR51E2. The y axis shows normalized AUC. (E) C3 concentration response at various fixed C7 concentrations in OR51E2. The y axis shows normalized AUC.

In WT, C7 is inactive (A). In contrast, C7 elicits robust activation in the pocket-expanded F155A and L158A variants, and these variants show reduced C3 responses relative to WT (B,C). C7 suppresses C3-evoked activation in a concentration-dependent manner across mixing schemes (D,E).

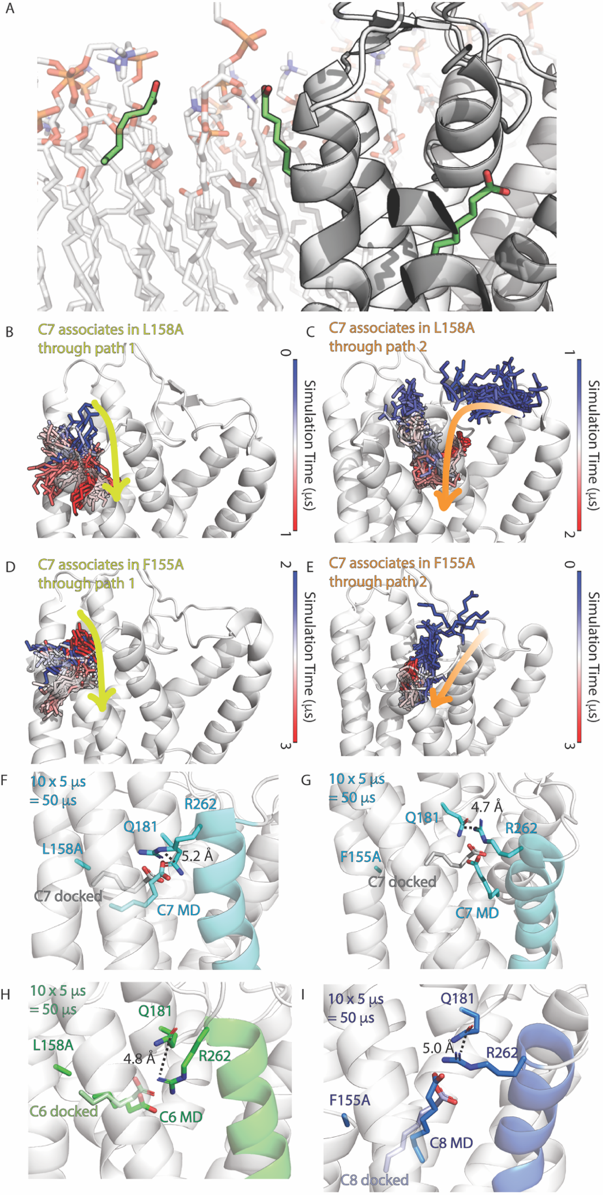

**Supplementary Figure 2. L158A, F155A mutants of OR51E2 allow productive binding of larger odorants, evident by ECL2-ECL3 closure.** (A) Illustration to show C7 insert its tail into membrane. The membrane lipids are drawn as white sticks. C7 are drawn as green sticks. OR structure is drawn as white new cartoon. (B, C) Path 1 and path 2 for C7 association in L158A as identified from MSM analysis of the MD simulation trajectories. (D, E) Path 1 and path 2 for C7 association in F155A. (F) Representative structure extracted MD simulations of the binding pose of C7 in OR51E2-L158A, docking pose of C7 is overlaid and distance between R262-Q181 is labeled. (G) Representative structure from MD simulations of the binding pose of C7 in OR51E2-F155A. (H) Representative structure from the association MD simulations of C6 in OR51E2-L158A overlaid with its docked pose. (I) Representative structure from the association MD simulations of C8 in OR51E2-F155A overlaid with its docked pose.

In both pocket-expanded mutants, C7 reaches a fully inserted bound pose and supports R262–Q181 gate closure, consistent with productive association (F,G). C6 and C8 also adopt docked-like bound poses in these variants while maintaining a closed gate (H,I).

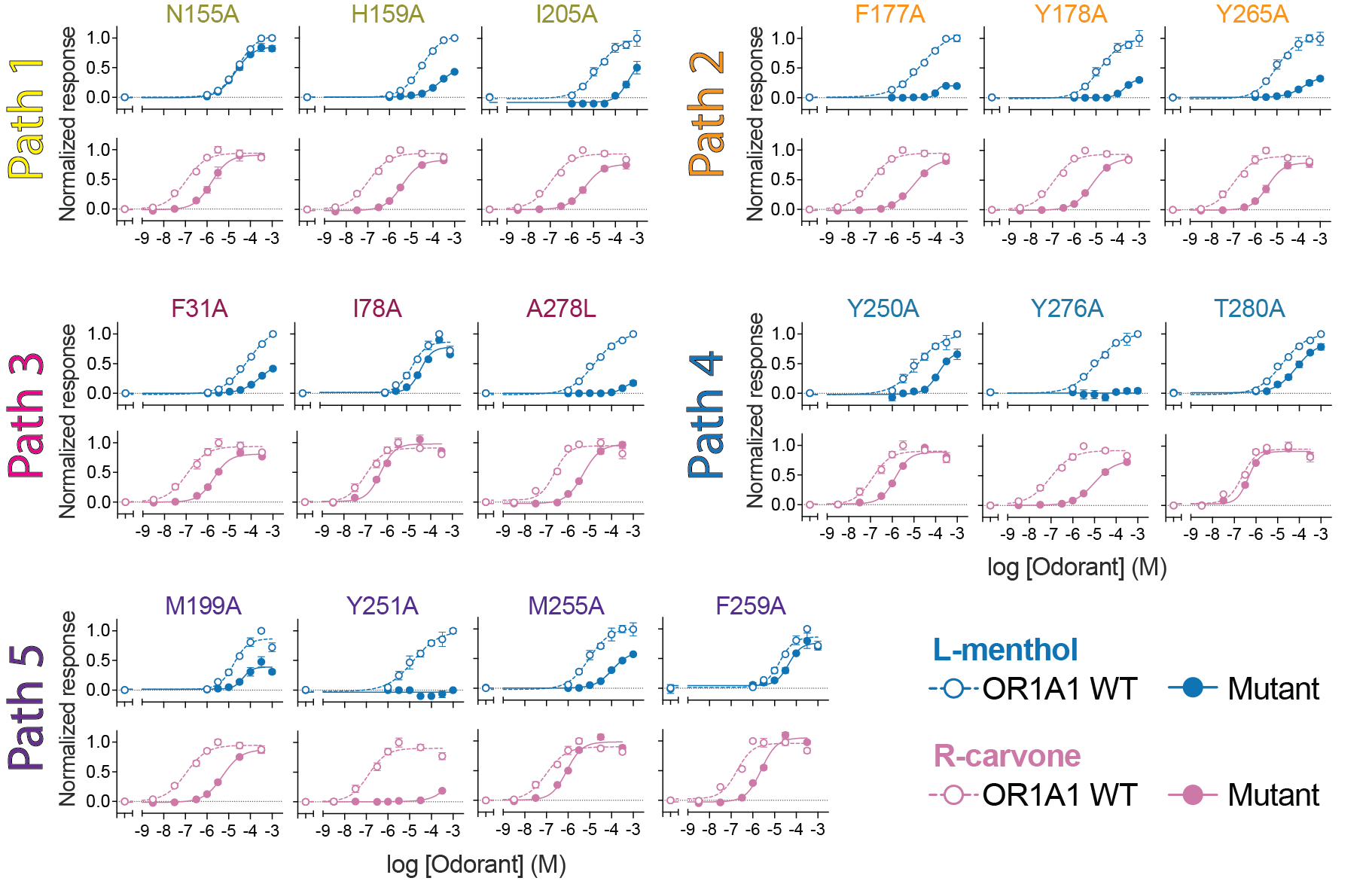

**Supplementary Figure 3. Mutational scan of OR1A1 gating residues along five association paths for L-menthol and R-carvone.** Each mini panel shows dose–response curves for OR1A1 WT and the indicated mutant. Blue curves show L-menthol responses, and pink curves show R-carvone responses. Open circles show OR1A1 WT, and filled circles show the mutant. The x axis shows log10 odorant concentration in molar units. The y axis shows the basal-subtracted GloSensor cAMP response normalized to the WT maximum response for the corresponding odorant. Path 1 includes N155A, H159A, and I205A. Path 2 includes F177A, Y178A, and Y265A. Path 3 includes F31A, I78A, and A278L. Path 4 includes Y250A, Y276A, and I280A. Path 5 includes M199A, Y251A, M255A, and F259A.

All mutants show potent or efficacy shift for at least one of the two odorants. Several gate mutations shift the relative responses to L-menthol versus R-carvone. These ligand-dependent effects are most prominent at gates along paths 1 and 5, including an R-carvone-biased effect at N155A and F259A and an L-menthol-biased effect at M255A, consistent with path-specific gating contributions to OR1A1 selectivity.

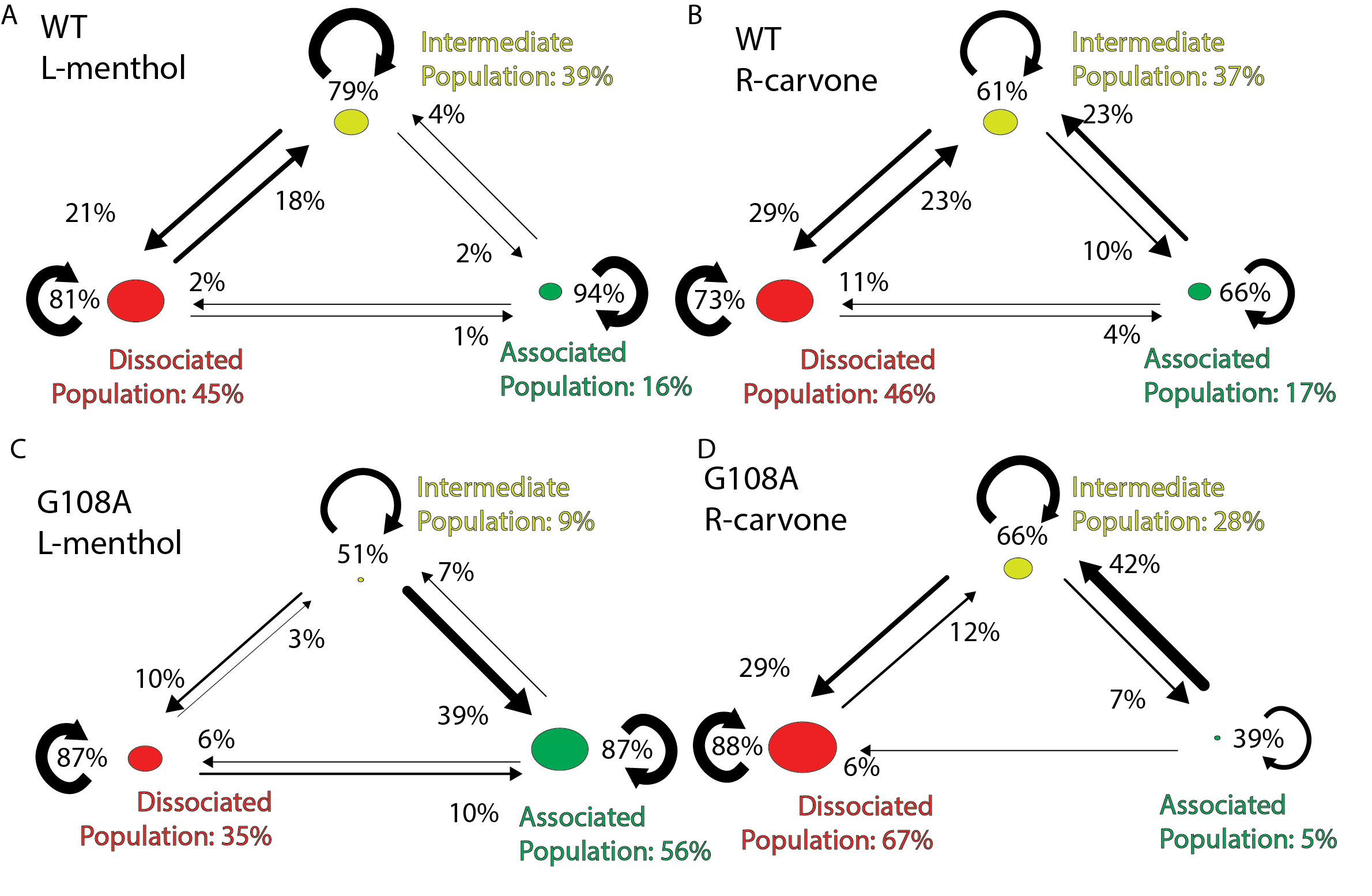

**Supplementary Figure 4. MSM-derived gating kinetics explain odorant-specific effects of the OR1A1 G108A variant.** Three-state Markov state models for (A) L-menthol WT, (B) R-carvone WT, (C) L-menthol G108A, (D) R-carvone G108A, showing transitions among Dissociated (red),  Intermediate (yellow), and Associated (green) macrostates. Node labels indicate populations for each macrostate, and arrows report MSM transition probabilities between or within states.

In WT, R-carvone shows a higher intermediate-to-associated transition probability than L-menthol, consistent with more frequent association under the sampled conditions (A,B). The G108A substitution reverses this kinetic bias by increasing the intermediate-to-associated transition probability for L-menthol while decreasing it for R-carvone (C,D), which matches the direction of the experimental potency shift.

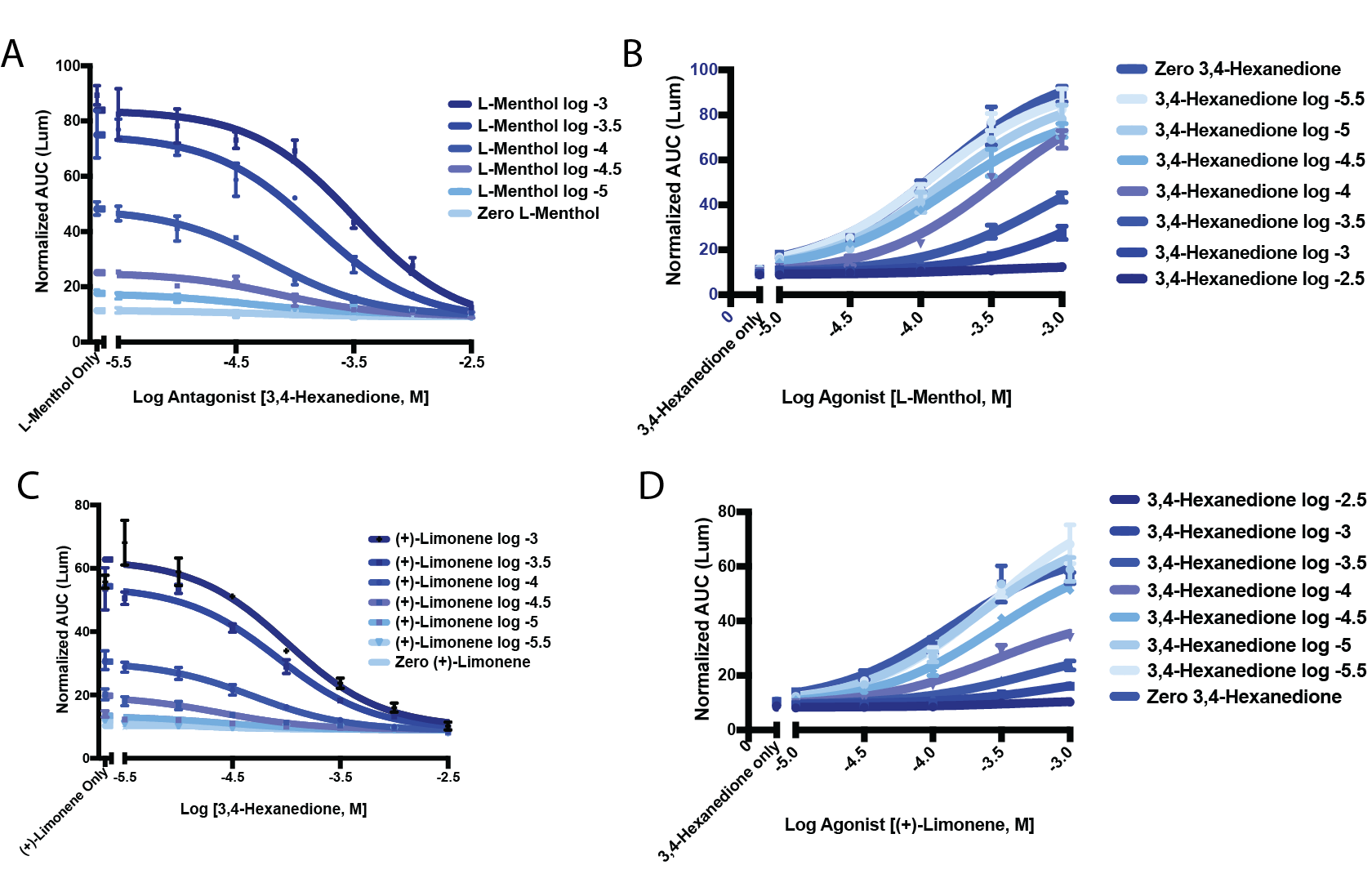

**Supplementary Figure 5. 3,4-hexanedione antagonizes OR1A1 activation by L-menthol and (+)-Limonene (**A) 3,4-hexanedione titration with fixed L-menthol concentrations. The y axis shows normalized GloSensor AUC. The x axis shows log10 3,4-hexanedione concentration in molar units. (B) L-menthol concentration response in the presence of increasing 3,4-hexanedione concentrations. The y axis shows normalized GloSensor AUC. The x axis shows log10 L-menthol concentration in molar units. (C) 3,4-hexanedione titration with fixed (+)-limonene concentrations. The y axis shows normalized GloSensor AUC. The x axis shows log10 3,4-hexanedione concentration in molar units. (D) (+)-Limonene concentration response in the presence of increasing 3,4-hexanedione concentrations. The y axis shows normalized GloSensor AUC. The x axis shows log10 (+)-limonene concentration in molar units. Points show the mean. Error bars show variability across replicates.

Increasing 3,4-hexanedione reduces responses elicited by both agonists and shifts the L-menthol and (+)-limonene dose–response curves downward, consistent with concentration-dependent antagonism (A–D).

**Extended Methods**

**Initial odorants position setup for both OR51E2 and OR1A1.**

We followed a published ligand-seeding protocol^1^ to construct a 3D odorant array on a 9×9×9 grid with a minimum pairwise spacing of x Å, where x was decided based on desired concentration. The array was rigidly aligned to the pre-equilibrated GPCR-membrane complex, after which odorants that sterically clashed with protein or lipids were removed (Supplementary Fig. 8A). To achieve the target solute concentration within the specified membrane/box dimensions, additional odorants were pruned from the farthest to the nearest relative to the receptor until the desired count was reached. The resulting GPCR-membrane-odorant assembly was then submitted to CHARMM-GUI^2–4^ for system building, solvation, neutralization, and parameter generation.

**Justification of Odorant Concentration and Lipids Composition.**

Odorant concentration and membrane lipid composition are critical parameters for interpreting ligand-association simulations of ORs. In explicit-solvent MD, ligand concentration is constrained by finite box size. In a typical simulation box with size around 100 × 100 × 110 Å, even a single odorant molecule corresponds to a nominal concentration in the 10^-3^ M range. Thus, 10^-6^ M range, such as EC50 concentration of L-menthol/R-carvone in OR1A1, cannot be reproduced with typical simulation box size. Consistent with other ligand-association MD studies, which commonly use ligand concentrations from approximately 10^-3^ to 10^-1^ M to enhance sampling,^5–7^ for example, Araki, et al. studied ligand association with a ligand concentration of 0.138 M, and justified their choice of high ligand concentration by stating that using many ligands increases the probability of capturing rare protein–ligand binding events within accessible MD timescales.^6^ We selected odorant concentrations by balancing experimental relevance with simulation stability. When possible, we used concentrations corresponding to experimentally active conditions, including EC50 or Emax concentrations, and further evaluated whether ligand association could be observed within accessible simulation timescales without destabilizing the GPCR fold or membrane environment. Based on these criteria, our final simulations used 0.18 M C3 (Emax) for OR51E2, 30 mM C6/C7/C8 (Emax) for OR51E2, and 8.54 mM L-menthol or R-carvone (Emax) for OR1A1. Detailed system-specific justifications for odorant concentration and lipid composition are provided below.

**Increasing Concentration of C3 with OR51E2 from EC50 to Emax concentration.**

We constructed an apo OR51E2 model from our recent cryo-EM structure by removing the G protein and the native odorant, propionic acid, and embedding the receptor in a mixed bilayer (75% POPC, 25% cholesterol).^8^ To probe spontaneous association, we added Emax concentration of C3 (0.18 M) to the bulk solvent, because preliminary trials at EC50 concentrations (9 mM) failed to yield binding events on accessible timescales (5 µs × 5 = 25 µs, Supplementary Fig. 6A, Supplementary Table 1). We then performed 75 µs of MD with this Emax concentration across five independent 15 µs replicas to capture the odorant association process.

**Choosing Emax Concentration of C7 with OR51E2.**

We construct C7 association system with Emax concentration of C7 in L158A/F155A mutants (30 mM), and ran 75 μs of MD (5 × 15 μs). We did not perform additional OR51E2-C7 simulations at the EC50 concentration because the primary objective was to compare C7 and C3 association under matched simulation conditions. Using comparable nominal odorant concentrations allowed us to evaluate whether the longer-chain odorant C7 altered the association pathway, gate occupancy, and receptor stability relative to C3, rather than introducing concentration as an additional variable.

**C3, C6, C7, C8 association simulation in OR51E2 L158A, F155A mutants at Emax concentrations.**

To efficiently examine whether C3, C6, C7, or C8 can associate into the odorant binding site of the OR51E2 L158A and F155A mutants, we initialized all systems from a single intermediate conformation extracted from the C7-WT association trajectory. In this intermediate conformation, R262 is oriented toward the solvent and there are two C7 molecules in the ECL2-ECL3 gate (Supplementary Fig. 8B). From this structure, we generated mutant systems by mutating L158 or F155 to alanine and replacing all C7 odorants with either C3, C6 or C8 at 30 mM concentration, yielding the set of systems summarized in Supplementary Table 1. Each system was then subjected to 10 independent production simulations of 5 µs. These simulations were designed to test whether odorants can associate from a partially bound intermediate state, rather than to construct MSM transition matrices.

**Reducing Concentration of L-menthol or R-carvone with OR1A1 and plasma membrane lipids composition from High concentration to Emax concentration.**

OR1A1 is a class II olfactory receptor whose native odorants are typically small, hydrophobic molecules.^9^ We sought to characterize how such odorants associate with class II ORs and how this contrasts with class I receptors, which preferentially bind water-soluble odorants. As an initial test, we generated an apo model from the consensus OR structure (PDB 8UXY) by removing the G protein and L-menthol, then embedded the receptor in a 75% POPC/25% cholesterol bilayer. The box was solvated with 0.18 M L-menthol, mirroring our OR51E2 setups. Across 60 µs of MD (5 × 12 µs), most L-menthol molecules partitioned into the membrane (Supplementary Fig. 6B), likely increasing lateral pressure at the GPCR–lipid interface and destabilizing the transmembrane bundle. Several trajectories showed helix separation and pronounced kinking, and no association events were observed (Supplement Table 1). This set of simulation suggest the odorant concentration need to be reduced, and lipids environment need to be considered.

To reduce membrane pressure and increase physiological realism, we next simulated an OR1A1 homology model (built on the consensus OR1 template) with L-menthol or R-carvone at 0.04 M in a plasma-mimetic bilayer. To pre-equilibrate the lipids, one consOR1 model was positioned in a 350 × 350 Å² plasma membrane,^10,11^ coarse-grained (CG) with Martini22p^12^ and simulated for 30 µs. From the final frame, a 100 × 100 Å² lipids patch around consOR1 was extracted and converted to all-atom. The OR1A1 model was then embedded in this all-atom environment, odorants were added into solution with same protocol as C3 in OR51E2, MD simulation models were prepared, and all-atom MD simulations were performed (5 × 15 µs; a total of 75 µs per system). Under these conditions, L-menthol associated with OR1A1 whereas R-carvone did not. However, receptor destabilization persisted, indicating that the concentration remained too high to preserve GPCR integrity over these timescales (Supplement Table 1) thus the concentration needs to be further reduced.

Guided by these outcomes, we further reduced the odorant concentration to Emax concentration (8.54 mM which includes 5 ligands in simulation box. EC50 concentration of 10^-5^ M or 10^-6^ M is not achievable due to simulation box size limitation), while maintaining the same lipid environment, aiming to minimize odorant-induced membrane pressure yet retain the possibility of observing spontaneous association (Supplement Table 1).

**Extra enriched MD simulations for sampling the five association paths in OR1A1.**

Because we determined that OR1A1 odorants mostly associate through the membrane, and that lipid composition and initial odorant placement may potentially bias association events, we sought to randomize lipid positions and distribute odorants near each TM region. Our goal was to ensure that observed association events are not artifacts of a particular odorant starting position or pathway. We first generated an array of odorants (L-menthol or R-carvone) similar as in the C3-OR51E2 system, then removed all odorants located in bulk solution or more than 8 Å from the GPCR (Supplementary Fig. 8C). This configuration was submitted to CHARMM-GUI to embed the receptor/odorant system into the plasma membrane-mimetic bilayer with the same lipid composition described in the Methods section. We performed 10 independent membrane-building runs, yielding 10 simulation systems with identical lipid composition but distinct initial lipid arrangements. Five systems were simulated for 5 µs and the remaining five for 2 µs, as the first 5-µs trajectories already exhibited odorant association on a reasonable timescale. These enriched simulations were designed to more fully sample the previously identified association pathways.

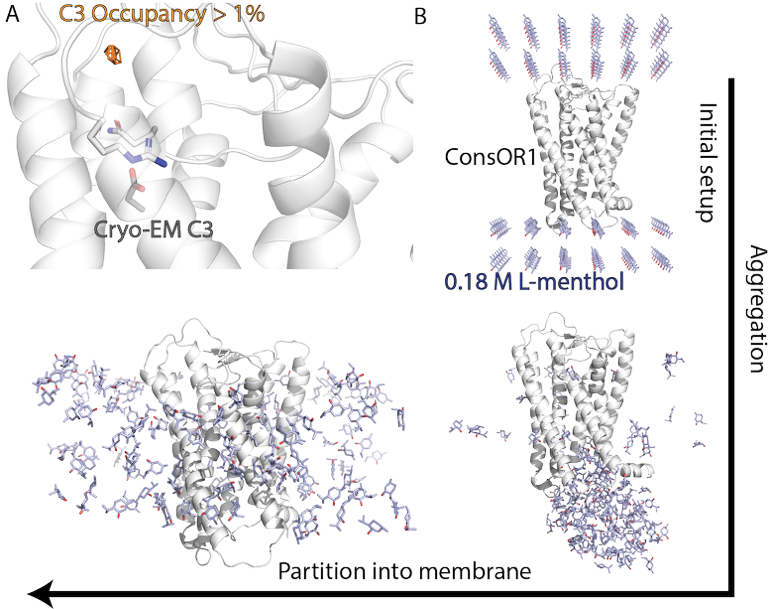

**Supplementary Figure 6. Odorant concentration tests that failed to yield binding events.** (A) C3 occupancy map for OR51E2 with odorants at EC50 concentration, highlighting regions with odorant occupancy ≥1% of trajectory time. (B) Initial 0.18 M L-menthol setup with the consOR1, odorants first cluster near the cytoplasmic G-protein–binding cavity and subsequently partition into the membrane.

**Robustness Consideration in MSM Construction**

To assess the robustness of our MSM analysis, we tested two modeling choices: (i) macrostate granularity and (ii) the contact distance cutoff used to construct the 2D distance array. In OR51E2, our baseline used four macrostates to differentiate the two association pathways. We rebuilt the MSMs using a coarser three-macrostates partition (associated, intermediate, dissociated) and varied the distance cutoff from 4.5 Å (a commonly used van der Waals contact threshold) to 6.5 and 8.5 Å. For both C3 and C7 association, the qualitative outcome was unchanged (Supplementary Fig. 7F): the C7 pathway consistently exhibits greater occupancy of intermediate states than C3. We therefore conclude that our method and result are robust to reasonable choices of macrostate definition and distance cutoff.

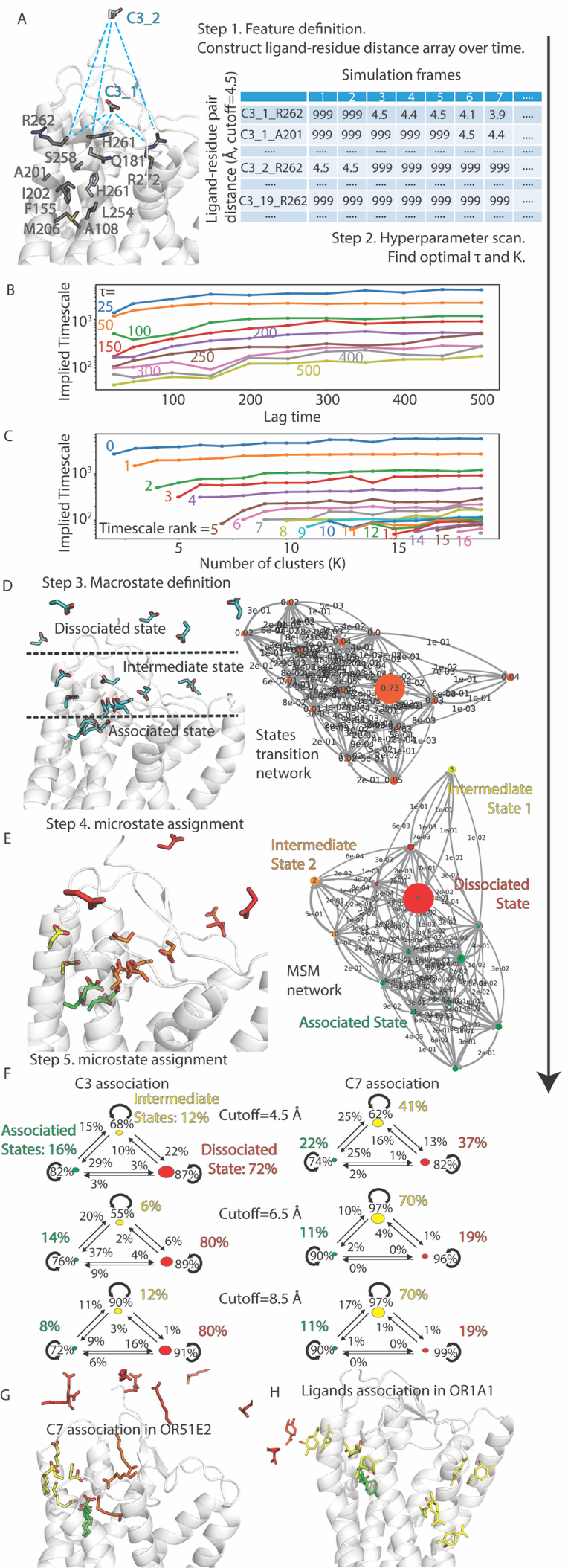

**Supplementary Figure 7. MSM validation and state exemplars.** (A) Schematic of the 2D distance array used as MSM input. (B) Implied timescales as a function of MSM lag time across a range of tICA lag parameters. (C) Implied timescales versus MSM lag time for varying numbers of clusters, K. (D) State-transition network for the 14 microstates of the C3 association in OR51E2. Representative odorant conformations for each microstate are projected onto the receptor. Multiple odorants are shown for dissociated microstates to illustrate spatial dispersion. One odorant exemplar is shown for other microstates. (E) Reduced MSM of C3 association in OR51E2 after assigning 14 microstates to 4 macrostates. Odorant conformations are colored by macrostate. (F) Reduced MSMs with 3 macrostates (associated, intermediate, dissociated) for C3 and C7 in OR51E2, evaluated for robustness under distance cutoffs of 4.5, 6.5, and 8.5 Å. (G) Representative C7 conformations from each of the 13 MSM microstates overlaid on OR51E2, colored by 4 macrostate. (H) Analogous representative conformations for R-carvone in WT OR1A1, colored by 3 macrostates.

**MSM Validation Supports Qualitative Macrostate Interpretation but Not Quantitative Kinetics**

To evaluate the reliability of the MSM-derived state assignments and population estimates, we performed additional validation analyses for both OR51E2 and OR1A1 odorant-association simulations. We assessed the reduced MSMs using 1) Chapman-Kolmogorov tests, 2) bootstrap and 3) jackknife resampling across independent trajectories, and 4) implied-timescale analyses over a range of lag times and clustering parameters. These analyses were performed for all the MD simulations in this study. The complete validation outputs, including summary metrics, population uncertainties, transition-probability uncertainties, CK-test plots, implied-timescale scans, and sensitivity analyses, are provided as Supplementary Data files.

The validation results showed that most reduced MSMs were suitable for qualitative macrostate interpretation, whereas a subset of systems showed finite-sampling uncertainty (Supplementary Data). For the better-supported systems, Chapman-Kolmogorov tests generally showed low-to-moderate mean errors of approximately 0.02~0.07, and bootstrap or jackknife analyses retained the dominant macrostate across trajectory resampling. For example, OR51E2 C3-F155A reproducibly favored the intermediate macrostate, with a baseline apparent occupancy of 0.94 and a bootstrap 95% confidence interval of 0.86~0.99, whereas OR51E2 C3-L158A reproducibly favored the dissociated macrostate, with a baseline apparent occupancy of 0.98 and a bootstrap 95% confidence interval of 0.97~0.99.

By contrast, only two systems out of all, OR51E2 WT-C3 and OR1A1 WT-L-menthol showed broader bootstrap uncertainty and stronger trajectory-to-trajectory variability, indicating that conclusions from these systems should be interpreted more cautiously. Across both systems, implied-timescale analyses did not show uniform plateau behavior over the tested lag-time ranges. We therefore use MSMs to support qualitative macrostate assignments and apparent occupancy trends, but not to derive quantitative transition probabilities, kinetic rates, or exact equilibrium populations.

We therefore interpret the MSMs primarily at the reduced macrostate level, where macrostates represent recurrent dissociated, intermediate, and associated-like odorant-position classes. These validation analyses indicate that the MSMs are appropriate for comparing qualitative apparent occupancy trends under matched simulation conditions, particularly when combined with direct trajectory inspection, odorant occupancy maps, residue-contact analyses, and functional mutagenesis data. However, the validation analyses also showed finite-sampling uncertainty, broad confidence intervals in some systems, and incomplete implied-timescale convergence. We therefore do not interpret MSM-derived populations as fully converged equilibrium populations, nor do we use MSM transition probabilities as quantitative kinetic rates. Instead, MSM results are used as supporting qualitative evidence for the proposed allosteric gating model, in which extracellular gate regions bias odorant association and selectivity.

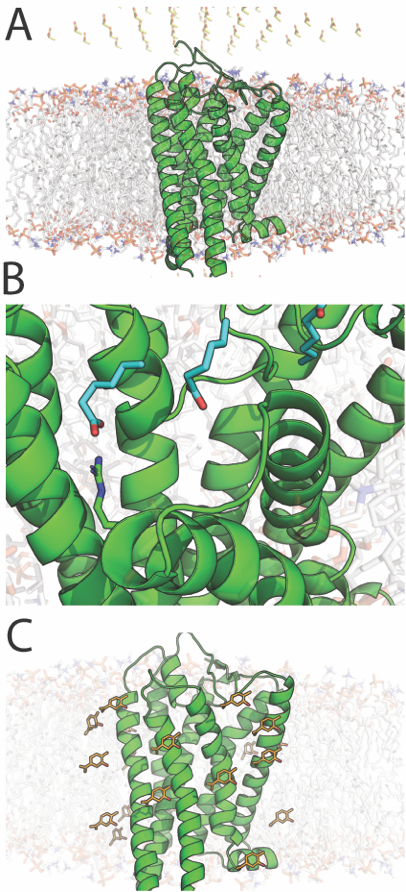

**Supplementary Figure 8. Different initial odorants position used in this study.** (A) Odorants were positioned into a array in solution, the number of odorants can be adjusted by the distances between array nodes. (B) The intermediate states from C7-WT simulation that was used to construct C3/C6/C7/C8 in F155A/L158A simulations. (C) Odorants positions in the OR1A1 enriched simulation.

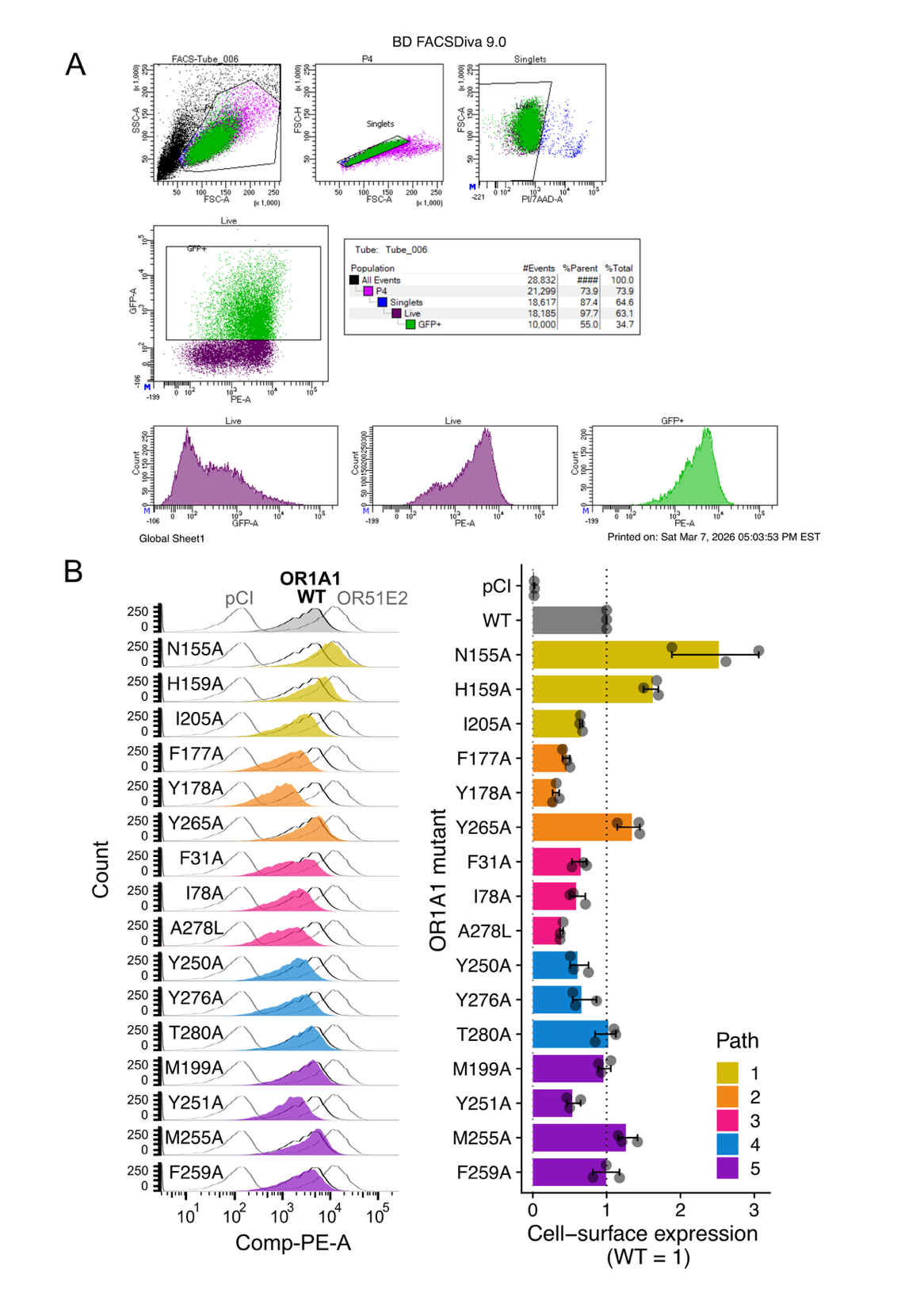

**Supplementary Figure 9. Flow cytometric analysis of cell-surface expression of OR1A1 mutants.**

**(A)** Representative gating strategy for analysis of cell-surface expression in HEK293T cells transfected with rho-tagged OR constructs together with RTP1S and eGFP. Cells were gated sequentially for the main population, singlets, live cells, and GFP-positive cells. Surface expression was measured by anti-rhodopsin staining followed by PE-conjugated secondary antibody detection. **(B)** Representative PE fluorescence histograms for OR1A1 WT and the indicated OR1A1 mutants, with pCI and OR51E2 shown as controls (left), and quantification of normalized cell-surface expression (WT = 1; right). Dots represent individual replicates; bars and error bars represent the mean and variability across replicates. Colors denote the path assignments shown in the key.

**Supplementary Table 1. Simulations performed in this study**

| GPCR | Odorants and  concentration | Initial odorant conformation | Simulation time | Lipids condition | Odorant association | GPCR deformation |
| --- | --- | --- | --- | --- | --- | --- |
| OR51E2  WT | C3  EC50 | Fig. S8A | 5 μs × 5  = 25 μs | 75% POPC  +  25% CHOL | No | No |
| OR51E2  WT | C3  Emax | Fig. S8A | 15 μs × 5  = 75 μs | 75% POPC  +  25% CHOL | Yes | No |
| OR51E2  WT | C7  Emax | Fig. S8A | 15 μs × 5  = 75 μs | 75% POPC  +  25% CHOL | Yes | No |
| OR51E2  WT | C3 Emax  C7 Emax | Fig. S8A | 12 μs × 5  = 60 μs | 75% POPC  +  25% CHOL | Yes | No |
| OR51E2  L158A | C6  Emax | Fig. S8B | 5 μs × 10  = 50 μs | 75% POPC  +  25% CHOL | Yes | No |
| OR51E2  F155A | C8  Emax | Fig. S8B | 5 μs × 10  = 50 μs | 75% POPC  +  25% CHOL | Yes | No |
| OR51E2  L158A | C7  Emax | Fig. S8B | 5 μs × 10  = 50 μs | 75% POPC  +  25% CHOL | Yes | No |
| OR51E2  F155A | C7  Emax | Fig. S8B | 5 μs × 10  = 50 μs | 75% POPC  +  25% CHOL | Yes | No |
| OR51E2  L158A | C3  Emax | Fig. S8B | 5 μs × 10  = 50 μs | 75% POPC  +  25% CHOL | No | No |
| OR51E2  F155A | C3  Emax | Fig. S8B | 5 μs × 10  = 50 μs | 75% POPC  +  25% CHOL | No | No |
| Cons OR1 | L-menthol  (0.18 M in solution) | Fig. S8A | 12 μs × 5  = 60 μs | 75% POPC  +  25% CHOL | No | Yes |
| OR1A1  WT | L-menthol  (0.04 M in solution) | Fig. S8A | 15 μs × 5  = 75 μs | Plasma membrane | Yes | Yes |
| OR1A1  WT | R-Carvone  (0.04 M in solution) | Fig. S8A | 15 μs × 5  = 75 μs | Plasma membrane | No | Yes |
| OR1A1  WT  Regular | L-menthol  (Emax in solution) | Fig. S8A | 15 μs × 5  = 75 μs | Plasma membrane | Yes | No |
| OR1A1  WT  Regular | R-carvone  (Emax in solution) | Fig. S8A | 15 μs × 5  = 75 μs | Plasma membrane | Yes | No |
| OR1A1  G108A  Regular | L-menthol  (Emax in solution) | Fig. S8A | 15 μs × 5  = 75 μs | Plasma membrane | Yes | No |
| OR1A1  G108A  Regular | R-carvone  (Emax in solution) | Fig. S8A | 15 μs × 5  = 75 μs | Plasma membrane | Yes | No |
| OR1A1  WT  Enriched | L-menthol  (Emax in membrane) | Fig. S8C | 5 μs × 5 +  2 μs × 5  = 35 μs | Plasma membrane | Yes | No |
| OR1A1  WT  Enriched | R-carvone  (Emax in membrane) | Fig. S8C | 5 μs × 5 +  2 μs × 5  = 35 μs | Plasma membrane | Yes | No |
| OR1A1  G108A  Enriched | L-menthol  (Emax in membrane) | Fig. S8C | 5 μs × 5 +  2 μs × 5  = 35 μs | Plasma membrane | Yes | No |
| OR1A1  G108A  Enriched | R-carvone  (Emax in membrane) | Fig. S8C | 5 μs × 5 +  2 μs × 5  = 35 μs | Plasma membrane | Yes | No |
| OR1A1  WT | 3,4-Hexiondione (Emax in solution) and  (+) Limonine (Emax in solution) | Fig. S8A | 15 μs × 5  = 75 μs | Plasma membrane | Yes | No |
|  |  |  | Total MD  length: 1260 μs |  |  |  |

**Supplementary Table 2. Association/Dissociation events observed in this study.**

| MD system | Velocity Index | Association Path and Event Index | Time step of contacting gate | Association time step | Lifetime at gate | Dissociation time step | Duration of association |
| --- | --- | --- | --- | --- | --- | --- | --- |
| OR51E2  WT with C3 | 1 | 1^st^ Event  Path1 | 1 μs | 1.2 μs | 0.2 μs | 4 μs | 2.8 μs |
|  | 2 | 1^st^ Event  Path 2 | 4.7 μs | 5.7 μs | 1.0 μs | Not observed | 9.3 μs |
|  | 2 | 2^nd^ Event  Path 2 | 7.1 μs | 7.3 μs | 0.2 μs | 9.6 μs | 2.3 μs |
| OR51E2  WT with C7 | 1 | 1^st^ Event  Path 1 | 6.8 μs | 7.6 μs | 0.8 μs | Not observed | 7.4 μs |
|  | 2 | 1^st^ Event  Path 1 | 0.6 μs | 6 μs | 5.4 μs | Not observed | 9 μs |
|  | 2 | 2^nd^ Event  Path 1 | 2 μs | 2.5 μs | 0.5 μs | 6.2 μs | 3.7 μs |
|  | 2 | 3^rd^ Event  Path 2 | 14.3 μs | 14.4 μs | 0.1 μs | Not observed | 0.6 μs |
|  | 3 | 1^st^ Event  Path 2 | 7.2 μs | 8.6 μs | 1.4 μs | Not observed | 6.4 μs |
|  | 3 | 2^nd^ Event  Path 2 | 7.4 μs | 7.6 μs | 0.2 μs | 7.8 μs | 0.2 μs |
| OR51E2  WT with C3 and C7 | 1 | C3 through Path 2  1^st^ Event | 2.2 μs | 2.3 μs | 0.1 μs | 5.3 μs | 2 μs |
|  | 1 | C7 through Path 2  2^nd^ Event | 10.5 μs | 10.6 μs | 0.1 μs | 11.4 μs | 0.9 μs |
|  | 2 | C7 through Path 2  3^rd^ Event | 11.2 μs | 11.21 μs | 0.01 μs | Not observed | 0.79 μs |
| OR51E2 L158A with C6 | 1 | Path 1  1^st^ Event | 2.6 μs | 2.9 μs |  | Not observed | 2.1 μs |
|  | 2 | Path 2  1^st^ Event | 0.4 μs | 0.6 μs |  | Not observed | 4.4 μs |
|  | 3 | Path 1  1^st^ Event | 0.8 μs | 3.1 μs |  | Not observed | 1.9 μs |
|  | 4 | Path 1  1^st^ Event | 0.2 μs | 0.6 μs |  | Not observed | 4.4 μs |
|  | 5 | Path 1  1^st^ Event | 0.6 μs | 0.8 μs |  | Not observed | 4.2 μs |
|  | 5 | Path 1  2^nd^ Event | 1.6 μs | 1.6 μs |  | Not observed | 3.4 μs |
|  | 6 | Path 1  1^st^ Event | 2.9 μs | 4.2 μs |  | Not observed | 0.8 μs |
|  | 7 | Path 1  1^st^ Event | 0.4 μs | 1 μs |  | 4.8 μs | 3.8 μs |
|  | 7 | Path 1  2^nd^ Event | 3.7 μs | 4.8 μs |  | Not observed | 0.2 μs |
|  | 8 | Path 1  1^st^ Event | 0.7 μs | 1.8 μs |  | Not observed | 3.2 μs |
| OR51E2 F155A with C8 | 1 | Path 1  1^st^ Event | 1.2 μs | 2.7 μs |  | 6.2 μs | 3.5 μs |
|  | 1 | Path 1  2^nd^ Event | 5.2 μs | 5.4 μs |  | 6.6 μs  through  Path 2 | 1.2 μs |
|  | 1 | Path 1  3^rd^ Event | 6.4 μs | 6.5 μs |  | 8 μs | 1.5 μs |
|  | 2 | Path 2  1^st^ Event | 5.8 μs | 5.9 μs |  | 6 μs | 0.1 μs |
|  | 3 | Path 1  1^st^ Event | 0.4 μs | 2.7 μs |  | 3.5 μs | 0.8 μs |
|  | 3 | Path 1  2^nd^ Event | 3.9 μs | 4 μs |  | 6.2 μs | 2.2 μs |
|  | 3 | Path 2  3^rd^ Event | 4.4 μs | 4.4 μs |  | 4.6 μs | 0.2 μs |
|  | 4 | Path 1  1^st^ Event | 0.7 μs | 1.5 μs |  | 8.6 μs | 7.1 μs |
|  | 4 | Path 2  2^nd^ Event | 7.9 μs | 8.6 μs |  | Not observed | 1.4 μs |
|  | 5 | Path 1  1^st^ Event | 0.5 μs | 2.8 μs |  | Not observed | 12.2 μs |
|  | 6 | Path 1  1^st^ Event | 1.2 μs | 2.7 μs |  | 2.8 μs | 0.1 μs |
|  | 6 | Path 2  2^nd^ Event | 2.7 μs | 2.8 μs |  | 4.4 μs | 1.6 μs |
|  | 7 | Path 1  1^st^ Event | 0.5 μs | 0.6 μs |  | Not observed | 9.4 μs |
|  | 8 | Path 1  1^st^ Event | 0.3 μs | 0.4 μs |  | Not observed | 9.6 μs |
| OR51E2 L158A with C7 | 1 | Path 1  1^st^ Event | 0.5 μs | 0.9 μs |  | 2 μs | 1.1 μs |
|  | 1 | Path 1  2^nd^ Event | 3.2 μs | 4μs |  | Not observed | 1 μs |
|  | 2 | Path 1  1^st^ Event | 0.5 μs | 0.9 μs |  | 1.8 μs | 0.9 μs |
|  | 2 | Path 1  2^nd^ Event | 3 μs | 4 μs |  | Not observed | 1 μs |
|  | 3 | Path 2  1^st^ Event | 0.6 μs | 0.8 μs |  | Not observed | 4.2 μs |
|  | 4 | Path 1  1^st^ Event | 0.8 μs | 2.3 μs |  | 3.2 μs | 0.9 μs |
|  | 4 | Path 1  2^nd^ Event | 2.6 μs | 4 μs |  | Not observed | 1 μs |
|  | 5 | Path 1  1^st^ Event | 0.1 μs | 1 μs |  | Not observed | 4 μs |
|  | 6 | Path 1  1^st^ Event | 0.1 μs | 0.5 μs |  | Not observed | 4.5 μs |
|  | 7 | Path 1  1^st^ Event | 0.1 μs | 0.4 μs |  | Not observed | 4.6 μs |
|  | 8 | Path 1  1^st^ Event | 2.2 μs | 2.3 μs |  | 3 μs | 0.7 μs |
|  | 8 | Path 1  2^nd^ Event | 3.2 μs | 3.3 μs |  | 3.5 μs | 0.2 μs |
| OR51E2 F155A with C7 | 1 | Path 1  1^st^ event | 1 μs | 4 μs |  | Not observed | 1 μs |
|  | 2 | Path 1  1^st^ Event | 0.9 μs | 2.6 μs |  | Not observed | 2.4 μs |
|  | 3 | Path 1  1^st^ Event | 3.7 μs | 4.4 μs |  | Not observed | 0.6 μs |
|  | 4 | Path 1  1^st^ Event | 0.8 μs | 1.2 μs |  | 2.8 μs | 1.6 μs |
|  | 5 | Path 2  1^st^ Event | 4.7 μs | 4.8 μs |  | Not observed | 0.2 μs |
|  | 6 | Path 1  1^st^ Event | 0.1 μs | 0.7 μs |  | Not observed | 4.3 μs |
|  | 7 | Path 1  1^st^ Event | 0 μs | 1.2 μs |  | 2 μs | 0.8 μs |
|  | 7 | Path 1  2^nd^ Event | 2.9 μs | 4.9 μs |  | Not observed | 0.1 μs |
| OR1A1  WT with L-menthol | 1 | Path 2  1^st^ Event | 0.9 μs | 2.8 μs | 1.9 μs | Not observed | 12.2 μs |
| OR1A1  WT with R-carvone | 1 | Path 5  1^st^ Event | 11.3 μs | 12 μs | 0.7 μs | 13.1 μs | 1.1 μs |
|  | 1 | Path 5  2^nd^ Event | 11.7 μs | 12 μs | 0.3 μs | 12.3 μs | 0.3 μs |
|  | 1 | Path 5  3^rd^ Event | 12 μs | 13.7 μs | 1.7 μs | Not observed | 1.3 μs |
|  | 2 | Path 5  1^st^ Event | 6.8 μs | 6.9 μs | 0.1 μs | 12.5 μs | 5.6 μs |
|  | 2 | Path 5  2^nd^ Event | 8.6 μs | 8.7 μs | 0.1 μs | 13.1 μs by Path 4 | 4.4 μs |
|  | 2 | Path 4  3^rd^ Event | 1.7 μs | 1.8 μs | 0.1 μs | 2.1 μs | 0.3 μs |
|  | 3 | Path 4  1^st^ Event | 14.2 μs | 14.9 μs | 0.7 μs | Not observed | 0.1 μs |
|  | 4 | Path 1  1^st^ Event | 2.1 μs | 2.2 μs | 0.1 μs | 3.7 μs | 1.5 μs |
|  | 4 | Path 3  2^nd^ Event | 14.5 μs | 14.6 μs | 0.1 μs | Not observed | 0.4 μs |
| OR1A1  G108A with L-menthol | 1 | Path 1  1^st^ Event | 1.7 μs | 2.1 μs | 0.4 μs | Not observed | 12.9 μs |
|  | 1 | Path 1  2^nd^ Event | 6.5 μs | 7.1 μs | 0.6 μs | 8.5 μs | 1.4 μs |
|  | 1 | Path 1  3^rd^ Event | 13.6 μs | 13.9 μs | 0.3 μs | Not observed | 1.1 μs |
|  | 2 | Path 4  1^st^ Event | 4.8 μs | 6 μs | 1.2 μs | 6.5 μs by path 1 | 0.5 μs |
|  | 2 | Path 1  2^nd^ Event | 6.5 μs | 6.8 μs | 0.3 μs | Not observed | 8.2 μs |
|  | 2 | Path 1  3^rd^ Event | 8.3 μs | 9.9 μs | 1.6 μs | 11.6 μs | 1.7 μs |
|  | 3 | Path 1  1^st^ Event | 1 μs | 1.3 μs | 0.3 μs | 2.2 μs | 0.9 μs |
|  |  | Path 4  2^nd^ Event | 5.4 μs | 5.5 μs | 0.1 μs | Not observed | 9.5 μs |
|  | 4 | Path 2  1^st^ Event | 0.1 μs | 0.2 μs | 0.1 μs | Not observed | 14.8 μs |
| OR1A1  G108A with R-carvone | 1 | Path 4  1^st^ Event | 14.0 μs | 14.1 μs | 0.1 μs | Not observed | 0.9 μs |
|  | 2 | Path 1  1^st^ Event | 14.8 μs | 14.8 μs | 0.02 μs | 14.9 μs | 0.1 μs |
| OR1A1  WT with L-menthol | 1 | Path 1  1^st^ Event | 3.5 μs | 4 μs | 0.5 μs | Not observed | 1 μs |
|  | 2 | Path 4  1^st^ Event | 1.2 μs | 2.1 μs | 0.9 μs | 2.1 μs | 0 μs |
|  | 3 | Path 5  1^st^ Event | 0 μs | 0.2 μs | 0.2 μs | Not observed | 4.8 μs |
|  | 3 | Path 5  2^nd^ Event | 0.1 μs | 3.6 μs | 3.3 μs | Not observed | 1.4 μs |
|  | 4 | Path 3  1^st^ Event | 0 μs | 0.4 μs | 0.4 μs | Not observed | 1.6 μs |
| OR1A1  WT with R-Carvone | 1 | Path 4  1^st^ Event | 0.17 μs | 0.18 μs | 0.01 μs | Not observed | 4.8 μs |
|  | 2 | Path 5  1^st^ Event | 0.1 μs | 0.2 μs | 0.1 μs | Not observed | 4.8 μs |
|  | 2 | Path 1  2^nd^ Event | 0.1 μs | 0.2 μs | 0.1 μs | Not observed | 4.8 μs |
|  | 3 | Path 5  1^st^ Event | 0.16 μs | 0.92 μs | 0.76 μs | Not observed | 1.08 μs |
|  | 4 | Path 1  1^st^ Event | 0 μs | 0.1 μs | 0.1 μs | Not observed | 1.9 μs |
| OR1A1  G108A with L-menthol | 1 | Path 5  1^st^ Event | 0.2 μs | 0.22 μs | 0.02 μs | Not observed | 1.78 μs |
| OR1A1  G108A with R-Carvone | 1 | Path 1  1^st^ Event | 0.6 μs | 0.67 μs | 0.07 μs | 1.3 μs | 0.63 μs |
|  | 2 | Path 5  1^st^ Event | 0.01 μs | 0.04 μs | 0.03 μs | Not observed | 2 μs |
|  | 3 | Path 1  1^st^ Event | 1.6 μs | 1.8 μs | 0.2 μs | Not observed | 0.2 μs |
|  | 4 | Path 2  1^st^ Event | 1.68 μs | 1.76 μs | 0.08 μs | Not observed | 0.24 μs |

**Reference.**
